# Dlg1 regulates subcellular distribution of non-muscle myosin II during *Drosophila* germband extension

**DOI:** 10.1101/2022.08.29.505652

**Authors:** Melisa A. Fuentes, Hayley N. Piper, Bing He

## Abstract

Elongation of the body axis through convergent extension is a conserved developmental process that is mediated by cell intercalation. During convergent extension of the germband epithelium in *Drosophila* embryos, planar polarized activation of non-muscle myosin II (“myosin”) promotes cell intercalation by facilitating patterned remodeling of adherens junctions. Here, we report that loss of the basolateral determinant Dlg1 leads to defects in the subcellular distribution of myosin during germband extension, and consequently, impairs proper junctional remodeling and apical area maintenance during cell intercalation. In *dlg1* mutant embryos, ectopic accumulation of myosin is observed at the medioapical domain and along the lateral membrane, whereas junctional myosin is greatly reduced. Analogous myosin mis-localization patterns are observed upon knockdown of other basolateral determinants, Scrib and Lgl, but not the apical determinants. The function of Dlg1 in regulating the spatial distribution of myosin requires its intact SH3 and GUK domains and involves the Rho1 GEF Cyst, active Rho1 and Rok. We propose that Dlg1 facilitates correct junctional remodeling and prevents undesired apical area variation during cell intercalation by regulating the subcellular location of myosin activation.

## Introduction

Convergent extension in epithelia provides an important mechanism to mediate tissue extension during morphogenesis (Huebner and Wallingford, 2018). During convergent extension, an epithelial sheet narrows in one direction (“convergence”) and elongates in the orthogonal direction (“extension”). This mode of tissue remodeling underlies a variety of morphogenetic processes, such as body axis elongation in *Drosophila* (“germband extension”), notochord extension in ascidians, and neural tube closure in chick embryos (Bertet et al., 2004; Irvine and Wieschaus, 1994; Munro and Odell, 2002; Nishimura et al., 2012; Zallen and Wieschaus, 2004). Convergent extension is mediated by cell intercalation, a process in which neighboring cells change positions. Cell intercalation is controlled by the planar polarity systems that define the planar organization of epithelial tissues (Butler and Wallingford, 2017; Devenport, 2014). While cell rearrangements during cell intercalation predominantly occur at the planar plane of the epithelia, studies in the past have revealed mechanisms that involve both the apical and basolateral regions of epithelial cells (Huebner and Wallingford, 2018). Our understanding of the interplay between apical-basal polarity and planar cell polarity and how it controls convergent extension is only beginning to emerge.

*Drosophila* germband extension is a well-studied example of convergent extension that occurs during early embryogenesis. It starts shortly after ventral furrow (VF) formation and near the beginning of posterior midgut (PMG) invagination (Kong et al., 2017; Paré and Zallen, 2020). During germband extension, the ectodermal tissue elongates and doubles in length along the anterior-posterior (AP) axis, by shortening along the dorsal-ventral (DV) axis. Both local and tissue-scale forces are necessary for germband extension. Local forces are largely driven by polarized cell intercalation. In the germband, two types of cell rearrangements occur during intercalation. At early stages, T1 transitions in cell quartets predominate. During a T1 transition, an anterior-posterior cell interface (AP junction) collapses, followed by the formation of a new, dorsal-ventral cell interface (DV junction) (Bertet et al., 2004). Over time, higher order rearrangements, like rosettes, become more prominent. During rosette formation, multiple AP junctions collapse such that neighbor exchange occurs within a group of 5-11 cells (Blankenship et al., 2006). Extrinsic forces that influence extension of the germband are primarily from the PMG, which is formed during gastrulation by invaginating the posteriorly localized endoderm primordium (Sweeton et al., 1991). PMG formation promotes tissue flow and dorsal-anterior movement of the germband (Butler et al., 2009; Collinet et al., 2015; Lye et al., 2015). In addition, pulling forces imposed on the germband by the PMG help orient and align the new junctions that are formed during cell intercalation (Collinet et al., 2015).

The molecular pathway that regulates germband extension has been extensively studied (Paré and Zallen, 2020). The pair-rule genes that are part of the anterior-posterior patterning system, *even-skipped* (*eve*) and *runt,* are required for planar cell polarity and polarized cell intercalation (Irvine and Wieschaus, 1994; Zallen and Wieschaus, 2004). Mutants that lack anterior-posterior patterning exhibit defective germband extension as a consequence of disrupted planar cell polarity and abnormal/reduced cell intercalation (Blankenship et al., 2006; Irvine and Wieschaus, 1994). Eve and Runt are transcription factors that activate expression of the Toll receptor proteins Toll-2, Toll-6, and Toll-8 in transverse stripes along the anterior-posterior axis of the embryo, which are essential for planar polarization of the germband epithelium (Paré et al., 2014). One of the crucial downstream events of Toll-2/6/8 expression is activation of non-muscle myosin II (hereafter “myosin”) in the germband, which provides the force necessary to shrink AP junctions during cell intercalation (Bertet et al., 2004; Blankenship et al., 2006; Paré et al., 2014). During germband extension, two populations of myosin can be observed at the apical region of the tissue. There is a pool of myosin that is enriched at AP junctions, and another pool of myosin that is localized to the medioapical cortex (Bertet et al., 2004; Rauzi et al., 2010; Zallen and Wieschaus, 2004). It has previously been suggested that pulses of medioapical actomyosin flow towards AP junctions helps promote shrinking, whereas junctionally enriched myosin functions to stabilize changes in AP junction length after shrinking.

Activation of myosin is mediated by the Rho1-Rok pathway. Rho1 is the *Drosophila* homologue of RhoA, which is a small GTPase that cycles between an inactive, GDP-bound form, and an active, GTP-bound form. The activation of Rho1 is mediated by its guanine nucleotide exchange factors (GEFs), which promote the exchange of GDP with GTP on Rho1. GTP bound Rho1 activates actomyosin contractility in a variety of physiological contexts by binding to and activating its downstream effectors Diaphanous, a formin family actin nucleator, and Rho-associated kinase (Rok) (Perez-Vale and Peifer, 2020). Diaphanous promotes actin assembly (Homem and Peifer, 2008; Johndrow et al., 2004), whereas activated Rok phosphorylates the myosin regulatory light chain, Spaghetti squash (Sqh), to initiate myosin activation and actomyosin network contraction (Winter et al., 2001). Activated Rok also inhibits the myosin regulatory light chain phosphatase, thereby further promoting myosin filament assembly and actomyosin network contraction (Amano et al., 2010). Like myosin, active Rho1 is enriched at AP junctions in a planar polarized manner in the germband tissue (Garcia De Las Bayonas et al., 2019). Active Rho1 is also observed at the medioapical cortex, where it activates polarized contractile flows of actomyosin towards vertical junctions (Garcia De Las Bayonas et al., 2019; Munjal et al., 2015). Two RhoGEFs, RhoGEF2 and Cyst/Dp114RhoGEF, have been reported to play important roles in regulating subcellular activation of Rho1 in the ectoderm during germband extension. RhoGEF2 activates Rho1 in the medioapical region of the cell, whereas Cyst primarily activates Rho1 at adherens junctions (Garcia De Las Bayonas et al., 2019; Silver et al., 2019). Knockdown of Cyst results in a loss of junctional myosin, and as expected, a lower number of T1 transitions and a slower rate of germband extension (Garcia De Las Bayonas et al., 2019). Cyst is uniformly distributed along adherens junctions, so it remains unclear how myosin activation is enriched at AP junctions (Garcia De Las Bayonas et al., 2019). Interestingly, a recent study has revealed that polarized actomyosin activation during germband extension involves interaction between Toll-8 and the adhesion GPCR Cirl, which may provide the upstream signal that activates Cyst in a planar polarized manner (Lavalou et al., 2021).

Several studies have shown that planar cell polarity and cell intercalation during germband extension are regulated by evolutionarily conserved protein complexes that determine the apical identity of epithelial cells. In polarized epithelia, the Crumbs(Crb)-PatJ-Stardust complex and Par3-aPKC-Par6 complex specify the apical domain, whereas the Scribble(Scrib)-Dlg1-Lgl complex specifies the basolateral domain (Rodriguez-Boulan and Macara, 2014; Tepass, 2012). In the planar polarized germband, the apical polarity protein, Bazooka (Baz)/Par3, along with adherens junctions, are enriched at DV cell interfaces. Furthermore, Baz is required for the proper enrichment of myosin at AP cell interfaces (Simões et al., 2010). Rho kinase (Rok), which is downstream of Toll-2/6/8 and is enriched at AP junctions, promotes enrichment of Baz at DV junctions by phosphorylating Baz and inhibiting its cortical localization at AP junctions (Simões et al., 2010). Despite being enriched at a complementary cortical domain, Baz may also function to promote myosin activation through Cyst. It has been recently shown that Cyst physically interacts with Baz and PatJ (Silver et al., 2019). In addition, recruitment of Cyst to the cortex is lost in *baz* and *crb* RNAi embryos at late stages of germband extension (Silver et al., 2019). Together, these results suggest that apical determinants regulate subcellular activation of myosin through multiple mechanisms during germband extension.

Despite the documented role of apical polarity determinants in regulating planar cell polarity and cell intercalation, the role of basolateral polarity determinants during germband extension has not been well characterized. It has been previously reported that *dlg1* null mutants exhibit abnormal segmentation in the epidermis at the end of germband extension, and subsequent germband retraction fails (Woods and Bryant, 1991). However, up until this point, it had remained unclear whether Dlg1 plays any role in elongation of the germband. We report that loss of Dlg1 results in severe myosin mis-localization defects during germband extension, including a reduction in junctional myosin, an increase in medioapical myosin, and ectopic accumulation of myosin along lateral membranes. This abnormal distribution of myosin is associated with aberrant T1 transitions and ectopic apical constriction in the ectodermal cells, which leads to defects in cell intercalation and tissue extension. These myosin mis-localization phenotypes are also observed in *scrib* and *lgl* RNAi embryos. The function of Dlg1 in regulating the spatial distribution of myosin is dependent on its SH3 and GUK domains and involves spatial regulation of Cyst by Dlg1. Together, our findings uncover a previously unappreciated role for Dlg1 in controlling the subcellular localization of myosin in the germband, and more importantly, for promoting cell intercalation and convergent extension.

## Results

### Loss of Dlg1 results in defects in tissue elongation shortly after the onset of germband extension

To determine if Dlg1 plays a role during germband extension, we first performed live imaging of the PMG in control and *dlg1* RNAi embryos. The effectiveness of the *dlg1* RNAi lines in knocking down Dlg1 has been previously demonstrated (Fuentes and He, 2022). During germband extension, the posterior midgut moves from the posterior end of the embryo towards the dorsal-anterior region of the embryo. The speed at which the PMG moves anteriorly is commonly used as a readout for the rate of germband extension (Garcia De Las Bayonas et al., 2019). We imaged embryos at the mid-sagittal plane and visualized the posterior midgut using the F-actin marker, UtrophinABD-Venus (Figard and Sokac, 2011), which marks the cell membrane (Fig. 1A). In *dlg1* RNAi embryos, PMG formation occurs like in the control embryos (Fig. 1A). However, the average rate of PMG movement away from the posterior pole is reduced in the *dlg1* RNAi embryos (Fig. 1B; control: 8.9 ± 1.6 μm/min, n = 8; *dlg1* RNAi: 5.6 ± 0.6 μm/min, n = 9; *p* = 0.0003). This result indicates that there is a significant reduction in the rate of germband extension in the *dlg1* RNAi embryos compared to the control embryos.

**Figure 1.**
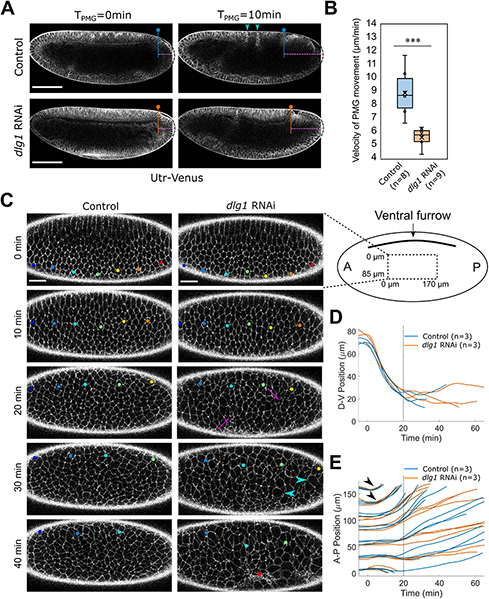
*dlg1* RNAi embryos exhibit a reduced rate of germband extension. **(A)** Movie stills showing the midsagittal plane of a representative control and *dlg1* RNAi embryo expressing UtrophinABD-Venus undergoing germband extension. The dorsal side is facing up. T_PMG_ = 0:00 (mm:ss) demarcates the onset of posterior midgut (PMG) formation. The position of the PMG in the control and the *dlg1* RNAi embryo is indicated by the blue and orange line, respectively. The magenta dotted line indicates the distance of the PMG away from the posterior end of the embryo. Arrowheads indicate dorsal folds, which fail to form in *dlg1* RNAi embryos. Scale bars: 100 μm. **(B)** Measurement of the rate of PMG movement between 60 and 140 μm from the posterior pole over in control and *dlg1* RNAi embryos. Unpaired, two-tailed student t-test was used for statistical analysis. ***: p <= 0.001. **(C)** Movie stills showing the subapical plane of a ventro-laterally oriented representative control and *dlg1* RNAi embryo expressing UtrophinABD-Venus during germband extension. The ventral side is facing up. The cartoon on the right depicts the position of the imaged region relative to the ventral furrow (VF). T = 0 minutes corresponds to the onset of rapid VF invagination. Colored dots indicate the individual cells that have been tracked over time. *Dlg1* RNAi embryos exhibit ectopic apical constriction after the onset of germband extension (magenta arrows in T = 20 minutes). Mitotic rounding can be observed in *dlg1* RNAi embryos as early as T = 30 minutes (cyan arrowheads) - approximately 20 minutes earlier than in the control embryos. The red star indicates the formation of an apical indentation associated with ectopic apical constriction in the *dlg1* RNAi embryos. Scale bars: 20 μm. **(D, E)** Quantification of the movement of cells along the DV (D) or AP (E) axis in control and *dlg1* RNAi embryos. Posterior movement of cells can be observed as early as T ≈ 5 minutes in both control and *dlg1* RNAi embryos (arrowheads). Dashed lines mark the time point when the rate of posterior movement starts to deviate between the control and *dlg1* RNAi embryos.

To further examine and compare the movement of the ectodermal cells in control and *dlg1* RNAi embryos, we performed live imaging on ventrolaterally oriented embryos expressing UtrophinABD-Venus during the first hour of germband extension. Then we tracked the displacement of cells initially located at different positions along the AP axis of the germband (Fig. 1C; Movie 1). We aligned embryos by the onset of rapid VF invagination, which we have defined as time zero for the following analyses. In both control and *dlg1* RNAi embryos, cell movement predominantly occurred along the DV axis during the first 10 minutes, as the VF invaginated (Fig. 1C, D). In control embryos, posterior movement of the germband cells began as early as T ≈ 5 – 10 min (Fig. 1E, arrowhead). Cells located closer to the posterior pole moved earlier, and the velocity of the movement was also higher compared to cells located more anteriorly (Fig. 1C, E). This graded pattern of tissue movement is expected for convergent extension and is consistent with previous measurements (Collinet et al., 2015). The pattern of AP tissue movement in the *dlg1* RNAi embryos was largely comparable to that of the control embryos during rapid VF invagination and during the initial phase of germband extension (T = 0 – 20 min; Fig. 1E). After 20 minutes, however, the rate of posterior movement slowed down in the *dlg1* RNAi embryos, in contrast to the more persistent movement observed in the control embryos (Fig. 1E). These observations confirm that germband extension is defective in *dlg1* RNAi embryos.

The impaired germband extension observed in the *dlg1* RNAi embryos is associated with aberrant apical cell shape changes in the ectodermal cells. During invagination of the VF, the morphology of the ectodermal cells in control and *dlg1* RNAi embryos was relatively comparable, and cell shape was largely uniform across the tissue in both genotypes (Fig. 1C, 0 – 10 min). During subsequent germband extension, however, the shape and size of the ectodermal cells became increasingly variable and heterogenous in the *dlg1* RNAi embryos (Fig. 1C, T = 20 min). Unlike the ectodermal cells in control embryos, many ectodermal cells in the *dlg1* RNAi embryos appeared to undergo apical constriction (Fig. 1C, magenta arrows). In some cases, apical constriction occurred in small patches of ectodermal cells and appeared to lead to the formation of ectopic shallow folds or pits (Fig. 1C, asterisk).

Interestingly, in addition to the tissue extension and apical cell morphology defects, the ectodermal cells in the *dlg1* RNAi embryos also displayed abnormal cell division patterns during germband extension. First, there appears to be an earlier onset of cell division in the *dlg1* RNAi embryos. In the region we examined in our experiments, cell division was first observed at T ≈ 50 minutes in control embryos (Movie 1). In the *dlg1* RNAi embryos, however, cell division could be detected as early as T ≈ 30 minutes (Fig. 1C, arrowhead; Fig. S1A). By T = 50 minutes, substantially more cells are dividing in the *dlg1* RNAi embryos than in the control embryos (Movie 1). The germband consists of multiple mitotic domains that enter mitosis at different times after cellularization (Foe, 1989). Since we oriented embryos ventrolaterally for imaging, we most likely captured cell division in mitotic domain N. However, the identity of this mitotic domain (particularly in the *dlg1* RNAi embryos) needs to be confirmed with future experiments. Second, we found that cell division in the *dlg1* RNAi embryos often incorrectly occurred along the vertical, apical-basal axis instead of along the horizontal, planar axis (Fig. S1B). This is consistent with previous studies that have reported a role for Dlg1 in regulating spindle orientation during cell division in multiple contexts (Bergstralh et al., 2013; di Pietro et al., 2016). It remains unclear how the defects in cell division affect the rate of germband extension in the *dlg1* RNAi embryos. Since the tissue in the *dlg1* RNAi embryos becomes progressively more disorganized during germband extension due to abnormal cell division, we decided to focus on characterizing the early germband extension defects prior to the occurrence of the abnormal cell division events.

### Germband cells in dlg1 RNAi embryos exhibit defects in junction remodeling during cell intercalation

Since cell intercalation plays an important role in tissue extension, we next asked if cell intercalation is defective in the *dlg1* RNAi embryos. To this end, we segmented sub-apical cell outlines and tracked cells over time. We focused our analysis on a 35-minute time window beginning at T ≈ 10 minutes. During this time window, the rate of germband extension began to deviate between the control and *dlg1* RNAi embryos. Cell tracking revealed that the extent of tissue extension was greatly reduced in the *dlg1* RNAi embryos compared to the control embryos (Fig. 2A). Importantly, no cell division was observed in the segmented tissue during this period. Therefore, the defects in tissue extension observed at this stage cannot be attributed to ectopic cell division in the mutant embryos.

**Figure 2.**
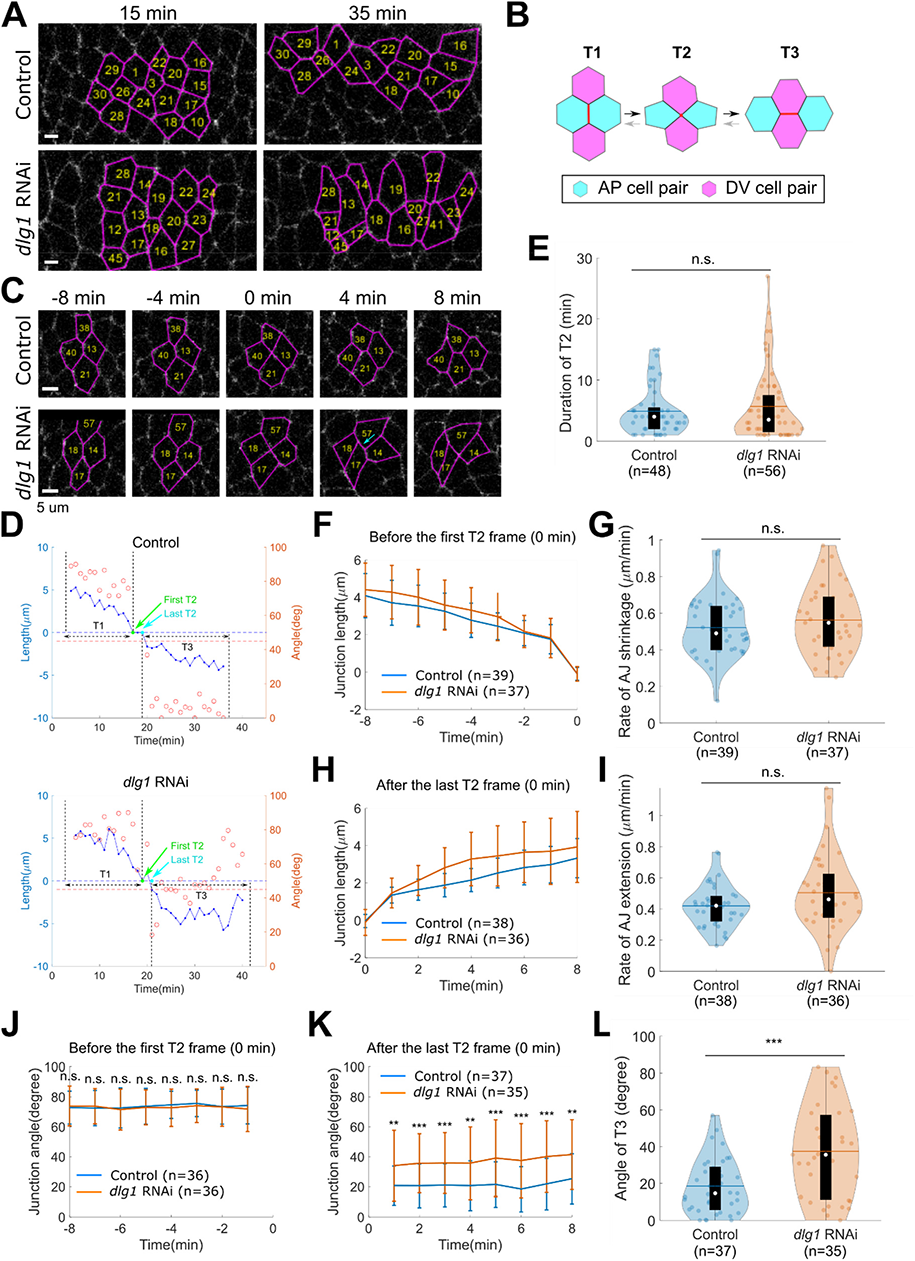
Germband cells in *dlg1 RNAi* embryos exhibit defects in junction remodeling during cell intercalation. **(A)** Extension of a group of cells in the germband tissue between T = 18 minutes and T= 37 minutes. The anterior side is on the left and the dorsal side is facing up (same below). Tissue extension along the AP axis is greatly reduced in the mutant compared to the control. Magenta: segmented cell outlines. Numbers indicate cell IDs. Scale bars: 5 μm. **(B)** Cartoon illustrating different quartet configurations during a T1 transition. During the T1-to-T2 transition, the AP junction (red vertical line) between the AP cell pair undergoes contraction and collapses into a vertex (red dot). During the T2-to-T3 transition, a new DV junction is formed (red horizontal line) between the DV cell pair. **(C)** Examples of individual control and *dlg1* RNAi quartets undergoing T1 transition. Arrow: oblique DV junction. Scale bars: 5 μm. **(D)** Plot illustrating the length and angle of AP and DV junctions over time. Data corresponds to the examples shown in C. Note that the negative sign was arbitrarily assigned to the length of DV junctions, which allowed us to simultaneously indicate the length and the type of the junctions. The angles range between 0° and 90°, with 0° indicating a horizontal junction (AP axis) and 90° indicating a vertical junction (DV axis). As exemplified in the plot, the DV junctions in the *dlg1* RNAi quartets often deviate from the AP axis. **(E)** Violin plot showing the distribution of the duration of T2, which is defined as the time between the first and last T2 configurations. Quartets from 3 control embryos and 4 *dlg1* RNAi embryos are shown (same below). **(F)** Average length of the AP junction over time before the first T2 configuration (defined as time 0 in the plot) in control and *dlg1* RNAi embryos. Error bars represent s.d. **(G)** Violin plot showing the average rate of AP junction shrinkage over an 8-minute time window before the first T2 configuration. **(H)** The average length of DV junctions over time in control and *dlg1* RNAi embryos following the last T2 configuration. Error bars represent s.d. **(I)** Violin plot showing the average rate of DV junction extension over an 8-minute time window after the last T2 configuration. **(J)** The average angle of AP junctions over time before the first T2 configuration. Error bars represent s.d. **(K)** Average angle of DV junctions over time after the last T2 configuration (defined as time 0 in the plot). Error bars represent s.d. **(L)** Violin plot showing the angle of DV junctions 6 minutes after the last T2 configuration. For all violin plots, the lower and upper limits of the black box correspond to the first and third quartiles (the 25th and 75th percentiles), the horizontal line indicates the mean, and the white dot in the black box indicates the median. Unpaired, two-tailed student t-test was used for all statistical analyses: n.s.: p > 0.05; *: p ≤ 0.05; **: p ≤ 0.01; ***: p ≤ 0.001 (unpaired, two-tailed student t-test).

During germband extension, groups of 4 cells, or cell quartets, undergo T1 transitions to promote cell intercalation (Fig. 2B). In control embryos, T1-transitions were frequently observed during the 35-minute time window that we examined. Consistent with previous reports, these T1-transitions were driven by shrinking of AP edges and extension of DV edges, which resulted in convergence of the tissue in the DV direction and extension of the tissue in the AP direction (Fig. 2C) (Bertet et al., 2004). Similarly, T1 transitions were also frequently observed in the *dlg1* RNAi embryos, indicating that loss of Dlg1 did not block T1 transitions (T2 counts per 30 min per 1000 μm^2^: 6.0 ± 0.7 in control embryos, n = 3 embryos; 4.9 ± 1.3 in *dlg1* RNAi embryos, n = 4 embryos; *p* = 0.63, Two-sided Mann-Whitney U-test). In order to better understand the behavior of cell quartets during T1 transitions, we examined individual quartets more closely. To this end, we analyzed 56 quartets from 3 control embryos and 63 quartets from 4 *dlg1* RNAi embryos. In both control and *dlg1* RNAi embryos, most quartets progressed through the T1:T2:T3 configurations in a chronological manner, undergoing stereotypical shrinking of AP junctions during T1 to T2 transitions and extension of DV junctions during T2 to T3 transitions (Fig. 2C, D). The duration between the first T2 configuration (the end of persistent AP junction shrinkage) and the last T2 configuration (the beginning of persistent DV junction extension) varied between quartets (Fig. 2E). In some cases when T2 duration was relatively long, cells underwent transient, temporary T2 to T1 and/or T3 to T2 reversals between the first and last T2 configurations (Fig. S2, arrows). The T2-to-T1 and T3-to-T2 reversals were observed in both control and *dlg1* RNAi embryos and did not appear to cause a difference in average T2 duration between the two genotypes (Fig. 2E; control: 4.9 ± 3.9 min; *dlg1* RNAi: 5.7 ± 5.8 min, mean ± std., *p* = 0.4, unpaired Student’s t-test).

To further explore the potential defects in junction remodeling in the *dlg1* RNAi embryos, we sought to compare the dynamics of AP junction shrinkage and DV junction extension between the control and the *dlg1* RNAi embryos. To this end, we measured junction length over time and determined 1) the rate of AP junction length change before the first T2 configuration (“rate of junction shrinkage”) and 2) the rate of DV junction length change after the last T2 configuration (“rate of junction extension”) (Fig. 2D). Unexpectedly, we found that the average rates of AP junction shrinkage and DV junction extension were indistinguishable between the control and *dlg1* RNAi embryos (Fig. 2F – I). The only noticeable mutant phenotype was a slight increase in the variation (as indicated by the standard deviation) of the junction extension rate (control: 0.42 ± 0.14 μm/min; *dlg1* RNAi: 0.50 ± 0.25 μm/min; Fig. 2I). These results indicate that loss of Dlg1 does not substantially affect the rate of junction shrinkage or the rate of junction extension during T1 transitions.

Interestingly, although the rate of new junction extension was comparable between the control and *dlg1* RNAi embryos, the orientation of the new junctions was abnormal in the mutant embryos. In the control embryos, the AP junctions remained largely vertical during T1-to-T2 transitions, with an average junction angle close to 70° (Fig. 2J). In contrast, the newly formed DV junctions that arose from T2-to-T3 transitions were predominantly oriented in the horizontal direction, with an average junction angle close to 20° (Fig. 2K). The average angle of the AP junctions in the *dlg1* RNAi embryos and that in the control embryos were comparable (Fig. 2J). However, the angle of the newly formed DV junctions in the mutant were much more variable (Fig. 2K, L). In some cases, the new DV junction resolved at a slant (Fig. 2C, arrow), and in more severe cases, the new DV junction resolved in the same direction as the old junction instead of in the orthogonal direction. The average DV junction angle was 37.4° ± 24.7° in *dlg1* RNAi embryos 6 minutes after the last T2 configuration, in contrast to 18.4° ± 15.0° in the control embryos (Fig. 2L). These observations reveal a prominent defect in the orientation of the newly formed DV junctions in the *dlg1* RNAi embryos. Since tissue extension mainly occurs during extension of the DV junctions (Collinet et al., 2015), we hypothesize that the new DV junctions that are formed at an oblique angle in the *dlg1* RNAi embryos are less effective at separating their AP cell pair. This hypothesis could well explain the reduced rate of tissue extension observed in the *dlg1* RNAi embryos.

### Germband cells in dlg1 RNAi embryos exhibit defects in apical cell area maintenance during cell intercalation

In addition to the DV junction orientation defects, another prominent defect that likely contributes to abnormal cell rearrangements in the *dlg1* RNAi embryos are the aberrant apical cell area changes in the germband. To further analyze this phenotype, we quantified the change in apical cell area in cell quartets over a 20-minute interval (ranging from T = 11 – 30 min to T = 21 – 40 min between different quartets). In control embryos, cell area generally remained constant during cell intercalation. In contrast, cell area in *dlg1* RNAi embryos was much more variable over time (Fig. 3A, B). Within the same quartet, some cells substantially reduced their apical area, while others apically expanded (Fig. 3B, C). In extreme cases, the apical domains of some cells constricted to such a small area that they were no longer clearly visible from the surface of the tissue (Fig. 3B, arrows). Although the area of the individual quartets was largely comparable between control and *dlg1* RNAi embryos (Fig. 3D), the area of cells within the same quartet in the *dlg1* RNAi embryos became much more heterogeneous at the end of the 20-min interval due to aberrant changes in apical area (Fig. 3E). Analysis of the net change in cell area during this 20-min interval (Δarea) further confirmed that the difference in Δarea among four cells within the same quartet was on average much greater in the *dlg1* RNAi embryos than in the control embryos (Fig. 3F). Furthermore, the mutant cells showed a greater temporal fluctuation in apical area (Fig. 3B, G; Methods). Together, these observations indicate that the size of the apical domain is less well maintained during germband extension in the absence of Dlg1.

**Figure 3.**
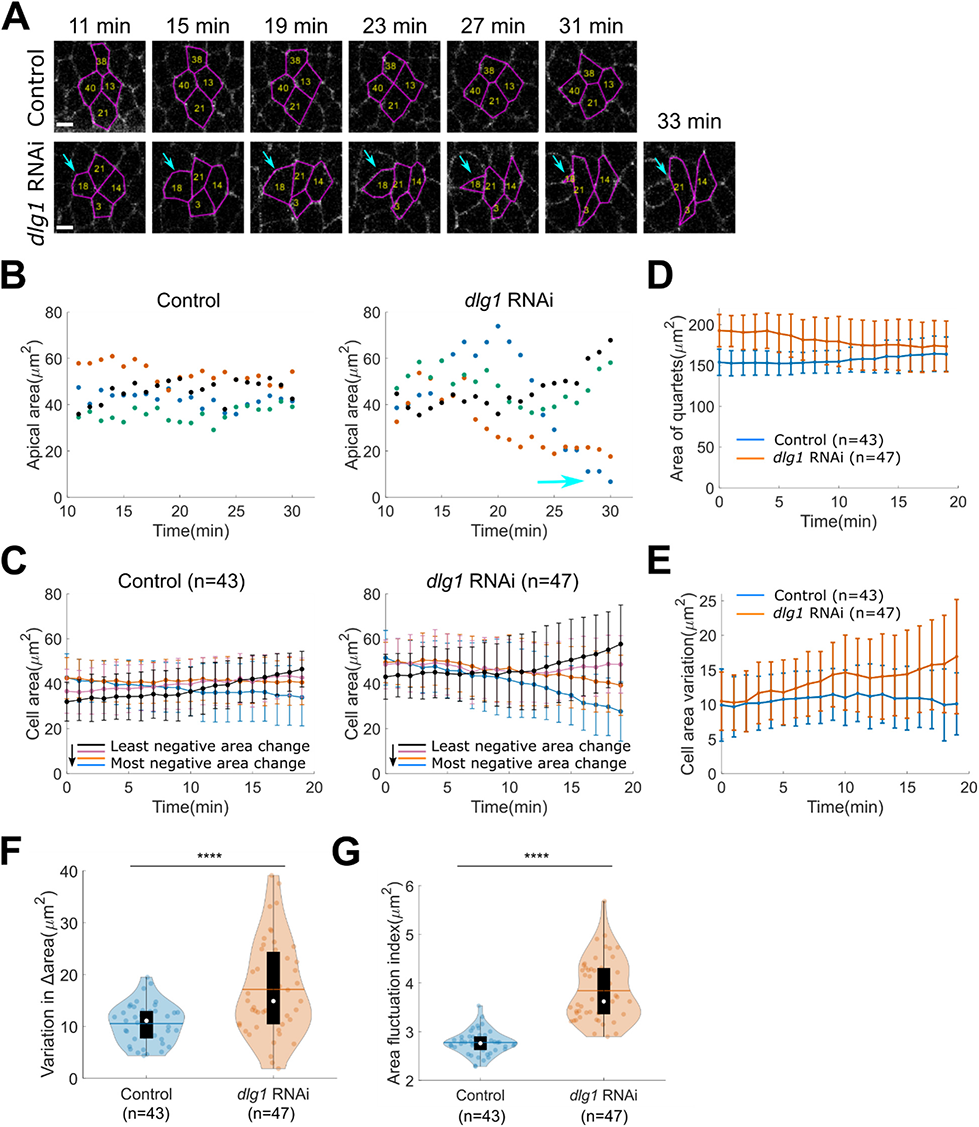
Germband cells in *dlg1 RNAi* embryos exhibit defects in apical cell area maintenance during cell intercalation. **(A)** Examples of individual control and *dlg1* RNAi quartets undergoing T transitions. Cyan arrows indicate a cell that undergoes ectopic apical constriction and eventually disappears from the surface. Magenta: segmented cell outlines. Numbers indicate cell IDs. Scale bars: 5 μm. **(B)** Comparison of apical cell area over time for cells in a control quartet and a *dlg1* RNAi quartet over time. T = 0 minutes indicates the onset of rapid VF invagination. Data corresponds to the examples shown in A. The cyan arrow indicates the cell denoted in A. **(C)** Cell area over time trend lines for four cells in the same quartet. Cells within a quartet were sorted into distinct categories based on the extent of negative area change over a 20-minute time interval (Methods). Cells from the same category are averaged between quartets of the same genotype. T = 0 minutes is the beginning of the 20-minute interval (same below). Cell area change significantly increased over time in *dlg1* RNAi embryos (47 quartets), but not in the control embryos (43 quartets). Error bars represent s.d. **(D)** Average apical area of individual quartets over time. Error bars represent s.d. **(E)** Comparison of the variation in cell area between four cells in the same quartet over time in control and *dlg1 RNAi* embryos. Error bars represent s.d. **(F)** Variation of area change over 20 minutes (Δarea) between four cells within the same quartet. **(G)** Index indicating the degree of temporal fluctuation of apical cell area for individual quartets (Methods). For all violin plots, the lower and upper limits of the black box correspond to the first and third quartiles (the 25th and 75th percentiles), the horizontal line indicates the mean, and the white dot in the black box indicates the median. Unpaired, two-tailed student t-test was used for all statistical analyses: ****: p ≤ 0.0001.

### Dlg1 promotes myosin localization to subapical adherens junctions in the ectoderm at early stages of germband extension

Next, we sought to understand the molecular defects underlying the junction remodeling and apical cell shape regulation phenotypes in the *dlg1* RNAi embryos. Planar polarization of myosin and myosin contractility at adherens junctions are important features that promote germband extension. We therefore wondered whether Dlg1 regulates the subcellular localization of myosin in the germband. To address this question, we performed live imaging of control and *dlg1* RNAi embryos expressing GFP-tagged Spaghetti Squash (Sqh), the myosin regulatory light chain, using a multiphoton microscope. In order to examine the apical/subapical plane of the embryo where adherens junctions are concentrated, we generated flattened *en face* projections at different apical-basal depths using a custom MATLAB script (Methods). After generating the flattened *en face* views, we examined Sqh-GFP signal in the lateral ectodermal cells adjacent to the constriction domain of VF. Approximately 4 minutes after the onset of VF invagination, we observed apical myosin accumulation in both control and *dlg1* RNAi embryos (Fig. 4A, “Subapical”, “T = 4 min”). This apical/subapical accumulation of myosin in the ectodermal cells increased over time. In control embryos, apical/subapical myosin was mostly localized to cell junctions. However, in *dlg1* RNAi embryos, apical/subapical myosin was rarely junctional, and instead, showed an increase in medioapical localization (Fig. 4A, magenta arrows). This ectopic, medioapical accumulation of myosin can be best appreciated by examining the most apical regions of the epithelium (Fig. S3, magenta arrows). This myosin phenotype is consistent with the observed ectopic apical constriction in the mutant tissue. To further evaluate the junctional localization of myosin in control and *dlg1* RNAi embryos, we defined a “junctional signal index” based on the repeated “peak-valley-peak” pattern of intensity distribution across multiple AP cell boundaries observed in wildtype embryos (Fig. S4A, B; Methods). Quantification confirmed that junctional myosin enrichment is lower in the lateral ectoderm of the *dlg1* RNAi embryos (Fig. 4B). Together, these results indicate that Dlg1 normally functions to promote myosin localization to apical adherens junctions and to inhibit medioapical accumulation of myosin.

**Figure 4.**
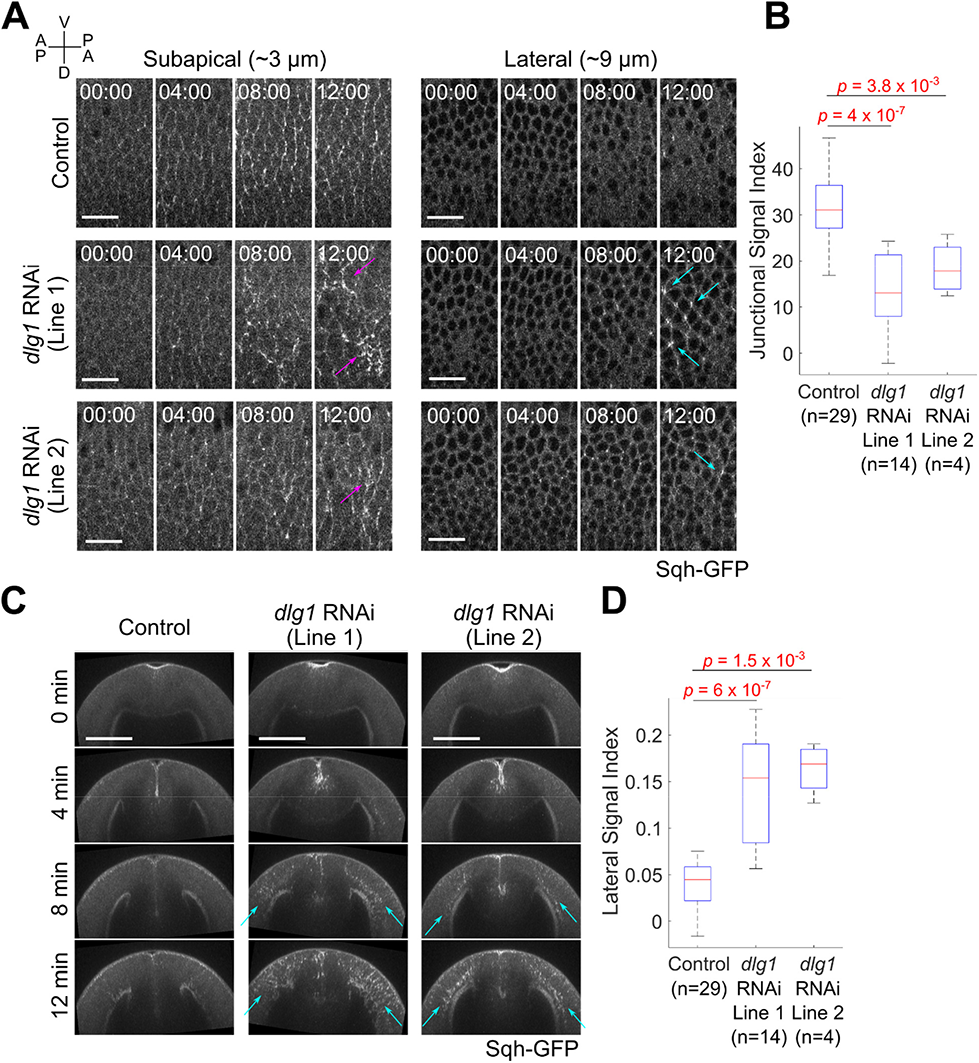
*dlg1* RNAi embryos exhibit myosin mis-localization defects in the ventrolateral ectoderm at early stages of germband extension. **(A)** Movie stills showing the flattened *en face* view of a representative control and two *dlg1* RNAi lines expressing the myosin marker, Sqh-GFP, at depths of 3 µm (subapical region, left panel) and 9 µm (lateral region, right panel) from the surface of the embryo. T = 0 minutes indicates the onset of VF invagination. Magenta arrows: ectopic medio-apical myosin. Cyan arrows: ectopic lateral myosin. Scale bars: 20 μm. **(B)** Junctional signal index of Sqh-GFP indicating the degree of myosin enrichment at the adherens junctions (Methods; Supplementary Figure 4). **(C)** Movie stills showing maximum projection cross-section views of the representative control and *dlg1* RNAi embryos shown in A. T = 0 minutes indicates the onset of VF invagination. Cyan arrows: ectopic lateral myosin. Scale bars: 50 μm. **(D)** Lateral signal index of Sqh-GFP indicating the degree of myosin enrichment at the lateral region of the germband epithelium (Methods; Supplementary Figure 4). Two-sided Mann-Whitney U-test was used for statistical comparison.

In addition to mis-localization of apical myosin, we found that germband cells in the *dlg1* RNAi embryos also exhibited ectopic myosin localization along their lateral membranes. To examine myosin localization along the apical-basal axis of the epithelium, we generated cross-section views of control and *dlg1* RNAi embryos and projected the maximum intensity of Sqh-GFP (Fig. 4C). In control embryos, myosin was strongly enriched at and tightly confined to the apical and basal ends of cells. Strikingly, in *dlg1* RNAi embryos, myosin lost its tight apical-basal confinement and decorated the lateral region of the cells (between the apical and basal membranes) (Fig. 4C, cyan arrows). Quantification of myosin signal intensity at the lateral region of the germband tissue confirmed this observation (Fig. 4D; Fig. S4C). To further characterize this phenotype, we generated flattened projections of control and *dlg1* RNAi embryos at ∼ 9 μm below the apical plane (Fig. 4A, “Lateral”). In control embryos, myosin was largely cytoplasmic in this region. In contrast, in *dlg1* RNAi embryos, myosin was localized to cell vertices and cell edges, with an apparent enrichment at cell vertices (Fig. 4A, cyan arrows). Aberrant accumulation of myosin in this region could be detected as early as T = 4 minutes and became more abundant over time. This finding suggests that in the ectoderm Dlg1 not only promotes junctional enrichment of apical myosin, but also inhibits localization of myosin at the lateral cell cortex. These myosin mis-localization phenotypes were observed in two different *dlg1* RNAi lines (Fig. 4), suggesting that they are not likely due to off target effects.

### dlg1 genetic mutant embryos exhibit similar defects in myosin organization as dlg1 RNAi embryos

To further confirm that Dlg1 plays a role in regulating the spatial organization of myosin, we examined Sqh-GFP in *dlg1* genetic mutants (Fig. 5A). For technical reasons, in all of our *dlg1* genetic mutant experiments, we used lines containing one copy of Sqh-GFP instead of two copies as in the RNAi experiments described above (Methods). In the *dlg1* RNAi background, reducing Sqh-GFP from two copies to one copy did not change the myosin mis-localization phenotype, although the intensity of the signal was reduced (Fig. S5). Since Dlg1 is largely maternally provided during early embryogenesis, as evidenced by the lack of an early phenotype in zygotic mutants (Perrimon, 1988), we first examined embryos derived from germline clones of a strong loss-of-function allele of *dlg1*, *dlg1^m52^*. These embryos are maternally mutant for *dlg1* but contain either zero copies or one copy of the wildtype *dlg1* gene from the paternal gamete (Methods). The molecular aberration in the *dlg1*^m52^ allele is a G to A point mutation which introduces a stop codon near the beginning of the third PDZ domain (after R482) and results in a severely truncated protein (Woods et al., 1996). Little to no protein can be detected by Western blot, suggesting that the *dlg1*^m52^ allele acts like a null (Woods et al., 1996). Similar to *dlg1* RNAi embryos, the *dlg1*^m52^ mutant embryos exhibited a decrease in junctional myosin and an increase in medioapical myosin (Fig. 5A, magenta arrows; Fig. 5B). In addition, myosin was mis-localized along the lateral membrane in the *dlg1^m52^* mutant embryos (Fig. 5A, C, cyan arrows; Fig. 5D).

**Figure 5.**
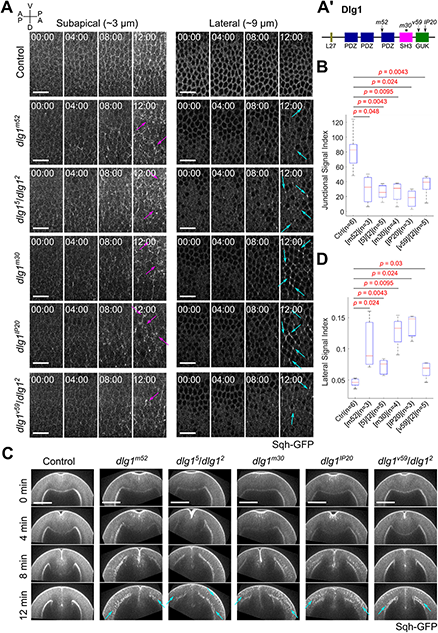
The SH3 and GUK domains of Dlg1 are both important for proper spatial organization of myosin in the ventrolateral ectoderm at early stages of germband extension. **(A)** Left: Schematic showing the domain organization of Dlg1 and the location of the mutations in the *dlg1* alleles used in this study. Asterisk: mis-sense mutation. Arrows: non-sense or frameshift mutations that result in premature stop codons. Right: Movie stills showing the flattened *en face* views of a representative control embryo and multiple representative *dlg1* mutant embryos expressing the myosin marker, Sqh-GFP, at depths of 3 µm (subapical region, left panel) and 9 µm (lateral region, right panel) from the surface of the embryo. T = 0 minutes indicates the onset of VF invagination. Magenta arrows: ectopic medioapical myosin. Cyan arrows: ectopic lateral myosin. Scale bars: 20 μm. **(B)** Junctional signal index of Sqh-GFP indicating the degree of myosin enrichment at the adherens junctions (Methods; Supplementary Figure 4). **(C)** Movie stills showing maximum projection cross-section views of the representative control and *dlg1* mutant embryos shown in A. T = 0 minutes indicates the onset of VF invagination. Cyan arrows: ectopic lateral myosin. Scale bars: 50 μm. **(D)** Lateral signal index of Sqh-GFP indicating the degree of myosin enrichment at the lateral region of the germband epithelium (Methods; Supplementary Figure 4). Two-sided Mann-Whitney U-test was used for statistical comparison.

Next, we examined a *dlg1* temperature sensitive allele. The *dlg1^2^* (*dlg1^H321F^*) mutant is associated with a 5.5 kb insert near the 5’ end of the gene (Woods and Bryant, 1991). *dlg1^2^* females are less fertile at 29°C compared to at 18°C, and the *dlg1^2^* allele does not produce any detectable protein at 29°C (Khoury and Bilder, 2020; Perrimon, 1988). We crossed *dlg1^2^* to another allele that is weakly temperature sensitive, *dlg1^5^* (*dlg1^lv55^*) (Perrimon, 1988), and examined embryos derived from the *dlg1^2^*/*dlg1^5^* mutant females. The *dlg1^2^*/*dlg1^5^* mutant embryos exhibited similar myosin mis-localization phenotypes as the *dlg1* RNAi embryos and the *dlg1^m52^* mutant embryos (Fig. 5). This result further confirms that Dlg1 regulates the spatial organization of myosin in the ectoderm during germband extension.

### The SH3 and GUK domains of Dlg1 are important for proper myosin localization

Dlg1 is a multi-domain protein with three PDZ domains, an SH3 domain, a HOOK domain, and a guanylate kinase (GUK) domain (Rodriguez-Boulan and Macara, 2014). In order to gain further insight into how Dlg1 regulates myosin, we examined available mutant alleles that are known to disrupt specific domains of Dlg1 (Fig. 5A’). The *dlg1^m30^* allele is marked by a T to C point mutation at base-pair 1895 in the SH3 domain, which changes the highly conserved leucine (L632) to a proline without affecting the expression level of the protein (Woods et al., 1996). The *dlg1^IP20^* allele is associated with a C to T point mutation at base-pair 2752 of the GUK domain, which converts a glutamine to a stop codon and eliminates the last third of the GUK domain (Woods et al., 1996). Likewise, the *dlg1^v59^* allele is a 14 base-pair deletion in the first third of the GUK domain that changes the open reading frame and introduces a stop codon after 12 amino acids, thereby eliminating the last two-thirds of the GUK domain (Woods and Bryant, 1991). Since *dlg1^v59^* is on a chromosomal inversion, we were not able to generate germline clones using the FRT/FLP technique. Instead, we crossed *dlg1^v59^* to *dlg1^2^* to reveal the phenotype of *dlg1^v59^*. Embryos derived from *dlg1^m30^* and *dlg1^IP20^* germline clones as well as the *dlg1^v59^*/*dlg1^2^* mutant embryos all displayed similar myosin mis-localization phenotypes as the strong loss-of-function *dlg1^m52^*allele (Fig. 5), suggesting that both the SH3 and GUK domains are important for the function of Dlg1 in mediating myosin organization in the germband.

### Scribble and Lgl also promote myosin localization to subapical adherens junctions in the ectoderm at early stages of germband extension

We next wondered if other apical-basal polarity proteins also regulate the spatial organization of myosin. To test this, we first knocked down the other two basolateral polarity determinants, Scribble (Scrib) and Lethal-giant larvae (Lgl), as described previously (Fuentes and He, 2022). Ectodermal cells in *scrib* RNAi and *lgl* RNAi embryos exhibited a similar loss of junctional myosin and increase in medioapical myosin, in addition to a similar increase in myosin along the lateral membrane (Fig. 6; Fig. S3). Furthermore, *scrib* and *lgl* RNAi embryos also displayed ectopic apical constriction and shallow pit/fold formation in the ectodermal tissue, as visualized by the F-actin marker, UtrophinABD-Venus (Fig. S6). These results suggest that basolateral determinants share a common function in regulating the subcellular localization of myosin in the germband.

**Figure 6.**
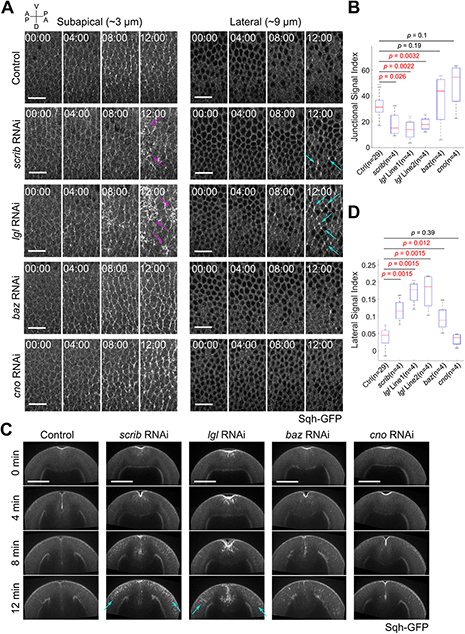
Knockdown of other basolateral polarity determinants, but not apical polarity determinants, results in myosin localization defects closely resembling those observed in the *dlg1* RNAi embryos. **(A)** Movie stills showing flattened *en face* views of representative control, *cno* RNAi, *baz* RNAi, *scrib* RNAi, and *lgl* RNAi embryos expressing the myosin marker, Sqh-GFP, at depths of 3 µm (subapical region, left panel) and 9 µm (lateral region, right panel) from the surface of the embryo. T = 0 minutes indicates the onset of VF invagination. Magenta arrows: ectopic medio-apical myosin. Cyan arrows: ectopic lateral myosin. Scale bars: 20 μm. **(B)** Junctional signal index of Sqh-GFP indicating the degree of myosin enrichment at the adherens junctions (Methods; Supplementary Figure 4). **(C)** Movie stills showing maximum projection cross-section views of representative control, *scrib* RNAi, *lgl* RNAi, *baz* RNAi, and *cno* RNAi embryos shown in A. T = 0 minutes indicates the onset of VF invagination. Cyan arrows: ectopic lateral myosin. Scale bars: 50 μm. **(D)** Lateral signal index of Sqh-GFP indicating the degree of myosin enrichment at the lateral region of the germband epithelium (Methods; Supplementary Figure 4). Two-sided Mann-Whitney U-test was used for statistical comparison.

To determine if the myosin phenotypes are specific to knockdown of the basolateral polarity determinants, we next decided to knockdown apical polarity determinants, such as Canoe (Cno) and Baz, and examine Sqh-GFP localization. Cno is the *Drosophila* homolog of Afadin and functions to physically link the actin cytoskeleton and adherens junctions (Sawyer et al., 2009). During apical-basal polarity establishment, Cno functions downstream of Rap1 and upstream of Baz, but can also be influenced by Baz (Bonello et al., 2018; Choi et al., 2013). Recently, Dlg1 and Scrib have also been reported to be able to influence the localization of Canoe (Bonello et al., 2019). To knockdown Cno, we used a previously described RNAi line that effectively reduces the levels of endogenous Cno to less than 5% during cellularization and gastrulation (Bonello et al., 2018). In *cno* RNAi embryos, normal recruitment of myosin to the adherens junctions was observed in the ectoderm, and myosin localized normally at apical and basal regions of the embryo (Fig. 6; Fig. S3). Thus, despite the important role of Cno in early polarity establishment prior to gastrulation, it is largely dispensable for the proper subcellular localization of myosin in the germband.

Next, we examined the role of the apical polarity determinant Baz, which is essential for formation of nascent subapical adherens junctions during cellularization (Harris and Peifer, 2004; Muller and Wieschaus, 1996). We used a previously described RNAi line to knockdown Baz (Takeda et al., 2018). *baz* RNAi embryos showed an intermediate phenotype. At the apical/subapical region, myosin was predominantly recruited to adherens junctions at early stages of germband extension, distinct from the *dlg1, scrib*, and *lgl* RNAi embryos (Fig. 6A; Fig. S3; T = 12 min). In some *baz* RNAi embryos, ectopic folds and/or pits formed in the lateral ectoderm, but only a moderate increase in medioapical myosin was observed in these regions (Fig. S7). Loss of Baz also caused a moderate accumulation of myosin along the lateral membrane, but this phenotype was not as severe as in the *dlg1*, *scrib* or *lgl* RNAi embryos (Fig. 6C, D). These results indicate that the apical determinant Baz is also involved in regulation of the subcellular localization of myosin in the germband. However, compared to the basolateral determinants, Baz seems to play a less important role in restricting myosin to the junctional and basal regions of cells.

Together, these observations suggest that regulation of myosin in the ectoderm by the basolateral determinants is at least partially independent of their role in regulating Cno and Baz localization. Furthermore, the lack of obvious myosin phenotypes in the *cno* RNAi embryos also highlights the complexity of the functional interactions between the apical/subapical polarity determinants during the course of polarity establishment.

### Dlg1 regulates localization of myosin activators Rok, active Rho1, and Cyst

Since myosin localization is abnormal in the *dlg1* RNAi embryos, we next wondered if the localization of myosin activators is also affected by knockdown of Dlg1. First, we examined localization of GFP-tagged Rho kinase (Rok-GFP) expressed from the *sqh* promoter (Abreu-Blanco et al., 2014) in control and *dlg1* RNAi embryos (Fig. 7A). In control embryos, Rok localized largely to the apical adherens junctions (Fig. 7A, arrowhead). Very little Rok signal was observed in deeper regions of the epithelium or along the lateral membrane (Fig. 7A, “Lateral”). This is consistent with myosin activation at adherens junctions (Bertet et al., 2004; Zallen and Wieschaus, 2004). Strikingly, Rok was significantly reduced and largely absent from the adherens junctions in the *dlg1* RNAi embryos (Fig. 7A, arrowhead; Fig. 7B). Ectopic medioapical accumulation of Rok could also be detected, albeit sparsely (Fig. 7A, magenta arrow). In addition, we occasionally observed an increase in Rok signal in lateral regions of the epithelium (Fig. 7A cyan arrows; Fig. S8A’). However, we could not draw a solid conclusion about this phenotype due to the relatively weak signal of Rok-GFP in deeper regions of the tissue (Fig. S8A, B). Together, this data suggests that the lack of myosin activation at the adherens junctions in the *dlg1* RNAi embryos is due to the loss of Rok from the junctions. It is also possible that ectopic myosin activation at the medioapical domain and along the lateral membrane is caused by ectopic accumulation of Rok, but more evidence is required to support this conclusion.

**Figure 7.**
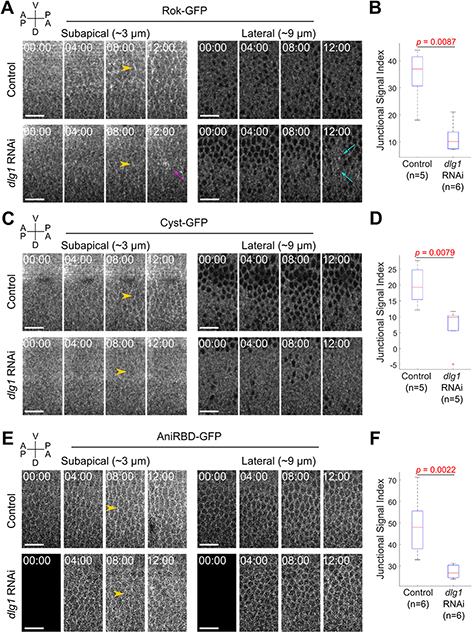
Knockdown of Dlg1 results in mis-localization of Rok, Cyst, and active Rho1 in the ventrolateral ectoderm at early stages of germband extension. **(A, C, E)** Movie stills showing the flattened *en face* views of a representative control and *dlg1* RNAi embryo expressing Rok-GFP (A), Cyst-GFP (C), and AniRBD-GFP (E) at depths of 3 µm (sub-apical region, left panel) and 9 µm (lateral region, right panel) from the surface of the embryo. A gaussian blur filter was applied to images shown in A and B. T = 0 minutes indicates the onset of VF invagination. The black images in (E) indicate data not acquired. Orange arrowheads: AP junctions. Magenta arrows: ectopic medio-apical Rok-GFP. Cyan arrows: ectopic lateral Rok-GFP. Scale bars: 20 μm. **(B, D, F)** Junctional signal index of Rok-GFP (B), Cyst-GFP (D) and AniRBD-GFP (F) indicating the degree of signal enrichment at the adherens junctions (Methods; Supplementary Figure 4). Two-sided Mann-Whitney U-test was used for statistical comparison.

Next, we decided to look further upstream the myosin activation pathway. In the *Drosophila* germband, the activation of Rho1, the activator of Rok, is regulated by the RhoGEFs, RhoGEF2 and Cyst/ Dp114RhoGEF. RhoGEF2 is known for its role in activating medioapical Rho1 in all three germ layers (Barrett et al., 1997; Hacker and Perrimon, 1998; Nikolaidou and Barrett, 2004; Dawes-Hoang et al., 2005; Garcia De Las Bayonas et al., 2019). Cyst, on the other hand, is expressed specifically in the ectoderm and promotes activation of Rho1 at adherens junctions during germband extension (Garcia De Las Bayonas et al., 2019; Silver et al., 2019). Since Cyst plays an important role in regulating junctional myosin in the germband, we hypothesized that Dlg1 promotes recruitment of myosin to apical adherens junctions by regulating the localization of Cyst. To test this hypothesis, we examined the localization of GFP-tagged Cyst (Cyst-GFP) expressed from the *sqh* promoter (Garcia De Las Bayonas et al., 2019) in control and *dlg1* RNAi embryos. Consistent with previous reports, Cyst was enriched at adherens junctions in control embryos (Fig. 7C) (Garcia De Las Bayonas et al., 2019; Silver et al., 2019). In contrast, in *dlg1* RNAi embryos, Cyst was absent from adherens junctions and was largely cytoplasmic (Fig. 7C, D). Unlike Sqh-GFP, we did not detect obvious ectopic localization of Cyst at the medioapical region or along the lateral membrane in *dlg1* RNAi embryos (Fig. 7C; Fig. S8C-D). However, because the signal of Cyst-GFP was relatively weak compared to Sqh-GFP, it remains to be determined whether the lack of prominent Cyst-GFP signal at the medioapical and lateral domains in the *dlg1* RNAi embryos is due to limited signal detection power in our experiments.

Consistent with the impaired junctional localization of Cyst-GFP, active Rho1, visualized by the Rho-binding domain of anillin (AniRBD-GFP) (Munjal et al., 2015), also showed reduced junctional localization in *dlg1* RNAi embryos (Fig. 7E, F). We were not able to evaluate the medioapical localization of AniRBD-GFP because of its relatively weak signal at the apical side compared to the autofluorescence of the vitelline membrane. Similar to Sqh-GFP, AniRBD-GFP also ectopically accumulated at the lateral membrane in *dlg1* RNAi embryos (Fig. S8E, F), suggesting that the lateral myosin phenotype might be caused by mislocalization of active Rho1. Together, our results suggest that the loss of junctional myosin in *dlg1* RNAi embryos is likely due to the lack of junctional recruitment of Cyst, which impairs the activation of myosin at the adherens junctions through the Rho1-Rok pathway.

In addition to Cyst, we also examined the localization of RhoGEF2 in control and *dlg1* RNAi embryos by using a RhoGEF2-GFP transgene expressed from the *sqh* promoter (Nakamura et al., 2017). In control embryos, RhoGEF2 was enriched at apical adherens junctions (Fig. S9A). Due to the bright autofluorescence signal from the vitelline membrane, we were not able to clearly visualize RhoGEF2 at the medioapical region of the germband cells as reported in a previous study (Garcia De Las Bayonas et al., 2019). In *dlg1* RNAi embryos, RhoGEF2 localized to apical adherens junctions, similar to the control embryos (Fig. S9A). Under our experimental conditions, no difference was observed between the control and the *dlg1* RNAi embryos at the medioapical domain or along the lateral membrane of the epithelium (Fig. S9A, B). Since RhoGEF2 still localizes to adherens junctions in the *dlg1* RNAi embryos, but junctional myosin is largely lost, it is unlikely that RhoGEF2 plays a major role in regulating junctional myosin, consistent with previous studies (Garcia De Las Bayonas et al., 2019).

### Ectopic activation of myosin along the lateral membrane in dlg1 RNAi embryos depends on the function of Cyst

Given the involvement of Cyst in Dlg1-mediated regulation of junctional myosin, we sought to determine whether Cyst is also involved in the ectopic activation of medioapical and lateral myosin in *dlg1* RNAi embryos. To this end, we examined myosin localization in *dlg1 cyst* double RNAi embryos and compared the phenotype to the single RNAi embryos. In *cyst* single RNAi embryos, a decrease in junctional myosin and a moderate increase in medioapical myosin was observed, which was consistent with previous reports (Fig. 8A, “Subapical”; Fig. 8B; Fig. S3) (Garcia De Las Bayonas et al., 2019; Silver et al., 2019). These phenotypes resembled the apical myosin phenotype in *dlg1* RNAi embryos, although the extent of medioapical myosin accumulation appeared to be milder in *cyst* RNAi embryos. On the other hand, in contrast to *dlg1* single RNAi embryos, myosin was properly segregated along the apical-basal axis and did not accumulate along the lateral membrane in *cyst* RNAi embryos (Fig. 8A, “Lateral”; Fig. 8C, D). In *dlg1 cyst* double RNAi embryos, a junctional myosin loss phenotype was observed, similar to both of the single RNAi mutants (Fig. 8A, “Subapical”; Fig. 8B). This observation is consistent with the view that Dlg1 and Cyst function in the same pathway in activating junctional myosin. Strikingly, in *dlg1 cyst* double RNAi embryos, little to no ectopic myosin was observed along the lateral membrane, suggesting that ectopic activation of lateral myosin in the *dlg1* RNAi embryos depends on the function of Cyst (Fig. 8A, “Lateral”; Fig. 8C, D). Interestingly, ectopic medioapical accumulation of myosin in the *dlg1 cyst* double RNAi embryos and the single *dlg1* RNAi embryos is largely comparable, suggesting that Cyst is dispensable for the activation of medioapical myosin in the *dlg1* RNAi embryos (Fig. 8A, “Subapical”; Fig. S3).

**Figure 8.**
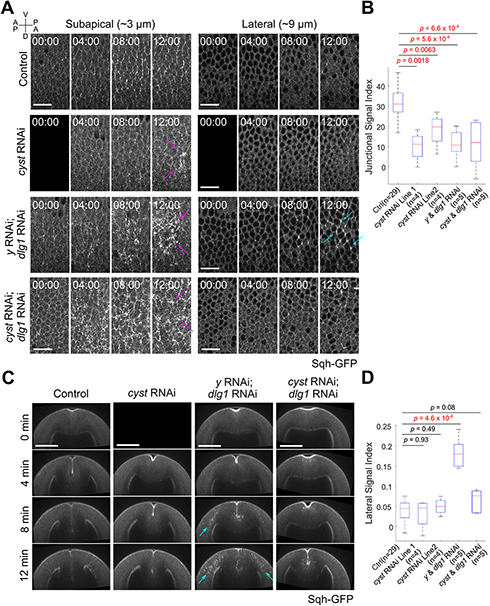
Knockdown of Cyst in the *dlg1* RNAi background rescues lateral myosin defects. **(A)** Movie stills showing the flattened *en face* views of representative control, *cyst* RNAi, *y dlg1* double RNAi, and *cyst dlg1* double RNAi embryos expressing the myosin marker, Sqh-GFP, at depths of 3 µm (sub-apical region, left panel) and 9 µm (lateral region, right panel) from the surface of the embryo. T = 0 minutes indicates the onset of VF invagination. The black images in (A) and (C) indicate data not acquired. Magenta arrows: ectopic medio-apical myosin. Scale bars: 20 μm. **(B)** Junctional signal index of Sqh-GFP indicating the degree of myosin enrichment at the adherens junctions (Methods; Supplementary Figure 4). **(C)** Movie stills showing maximum projection cross-section views of the representative control, *cyst* RNAi, *y dlg1* double RNAi, and *cyst dlg1* double RNAi embryos shown in A. T = 0 minutes indicates the onset of VF invagination. Scale bars: 50 μm. **(D)** Lateral signal index of Sqh-GFP indicating the degree of myosin enrichment at the lateral region of the germband epithelium (Methods; Supplementary Figure 4). Two-sided Mann-Whitney U-test was used for statistical comparison.

**Figure 9.**
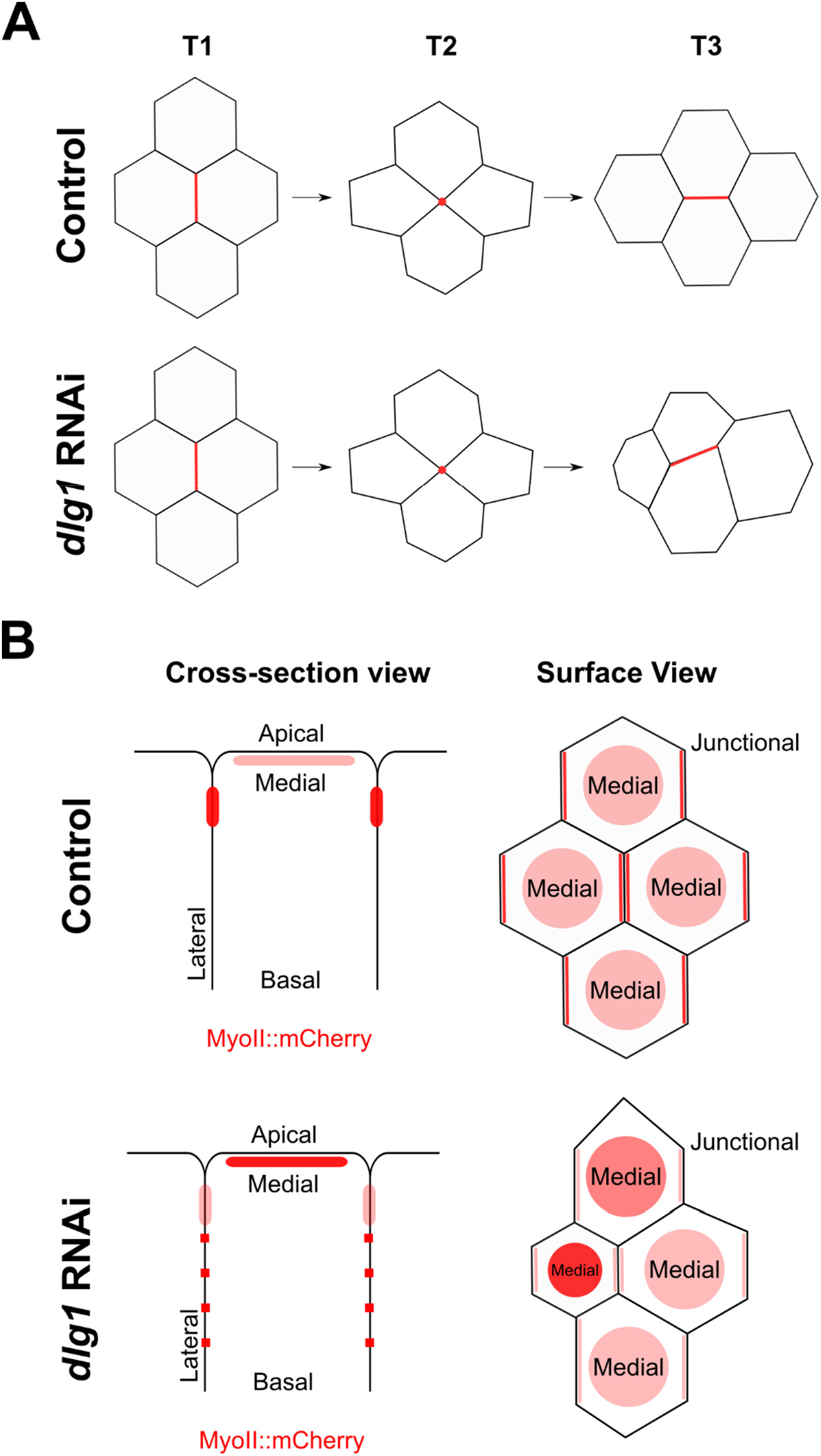
Dlg1 is essential for promoting proper myosin distribution and orchestrating cell rearrangements during germband extension. **(A)** During germband extension, cell intercalation is facilitated by T1 transitions. The AP cell interface shrinks along the D-V axis, resulting in T1-to-T2 transition. Subsequent resolution and extension of the new DV cell interface converts the T2 configuration into a T3 configuration. The new DV junctions are largely parallel to the A-P axis in the control embryos. Dlg1 regulates T1 transitions during germband extension. In the absence of Dlg1, T1 transitions often resolve and extend in a slant. In addition, maintenance of apical cell area is impaired in the *dlg1* RNAi embryos**. (B)** Dlg1 regulates myosin distribution in ectodermal cells during germband extension. In wildtype embryos, myosin is tightly confined to the apical and basal regions of the epithelium. Near the apical plane, myosin is medio-apically localized and planar polarized, such that it is enriched at junctions along the DV axis. In *dlg1* RNAi embryos, myosin loses its tight confinement to the apical and basal regions of the epithelium and instead decorates the lateral membrane. In addition, at the apical plane, medio-apical myosin levels are higher and junctional myosin levels are lower than in the control.

To rule out the possibility that the apparent rescue of the lateral myosin phenotypes in the *dlg1* RNAi embryos through knockdown of Cyst was due to a dilution effect that reduced knockdown efficiency in the double RNAi experiments, we examined myosin localization in *dlg1 yellow* double RNAi embryos. In this scenario, *yellow* RNAi served as a control because *yellow* does not have a known function in oogenesis or early embryogenesis (FlyBase). In *dlg1 yellow* double RNAi embryos, a decrease in junctional myosin, an increase in medioapical myosin, and an increase in lateral myosin were observed, much like in the single *dlg1* RNAi embryos (Fig. 8; Fig. 4). These results indicate that knockdown of Dlg1 in our double RNAi experiments was effective. Together, our results suggest that ectopic accumulation of myosin at the lateral domain in the *dlg1* RNAi embryos is attributed to mis-activation of Cyst upon loss of Dlg1 function.

## Discussion

In this work, we have found that the basolateral determinants Dlg1, Scrib, and Lgl play an important role in regulating the subcellular and spatial organization of myosin in the ventrolateral ectoderm during germband extension. Up to this point, the role of the basolateral determinants during germband extension had not been extensively characterized. We found that knockdown of any of the three basolateral determinants, Scrib, Dlg, or Lgl, resulted in decreased levels of junctional myosin and increased levels of medioapical myosin at the apical domain. Furthermore, loss of any of the three basolateral determinants resulted in ectopic myosin activation along the lateral membrane. These myosin phenotypes are also observed in *dlg1* genetic mutants. Knockdown of apical polarity determinants, Cno and Baz, on the other hand, did not recapitulate these phenotypes. Through structure function analysis, we found that the SH3 and GUK domains of Dlg1 are both important for the subcellular spatial organization of myosin in the ectoderm. Finally, we present evidence suggesting that Dlg1 regulates the spatial organization of myosin partially through regulation of the Rho1 GEF Cyst. We observed that in the ectoderm of *dlg1* RNAi embryos, Cyst, active Rho1, and Rok all became depleted from junctions. Furthermore, we observed that depletion of Cyst in *dlg1* RNAi embryos prevented ectopic accumulation of myosin along the lateral membrane, but not at the medioapical domain. Together, these data suggest that Dlg1 regulates the spatial organization of myosin during germband extension partly through the Cyst-Rho1-Rok pathway.

By analyzing the cellular defects in *dlg1* RNAi embryos during germband extension, we found that the defects in myosin localization were associated with abnormal junction remodeling and apical area regulation during cell intercalation. In particular, we observed that the orientation of the newly formed DV junctions was no longer constrained along the AP axis in the *dlg1* RNAi embryos. Additionally, apical cell area became much more heterogeneous over time in the mutant embryos. We frequently observed cells that underwent ectopic apical constriction. This caused other cells within the same quartet to expand their apical domain. We also observed ectopic accumulation of myosin at the medioapical domain in a subset of cells in the *dlg1* RNAi embryos, which likely caused the observed ectopic apical constriction. Based on these observations, we propose that Dlg1 facilitates effective convergent extension by regulating appropriate myosin localization in germband cells. Lacking functional Dlg1 results in reduction of junctional myosin and aberrant accumulation of medioapical myosin. This leads to ineffective T1 transitions and ectopic apical constriction, which ultimately leads to defects in tissue extension. In addition to the aforementioned defects, we also observed other phenotypes in the *dlg1* RNAi embryos during germband extension. In particular, we observed an earlier onset of cell division in the germband of *dlg1* RNAi embryos, and we noticed that the plane of cell division was often mis-oriented. *dlg1* RNAi embryos also exhibited defects in the formation of the transverse folds at the dorsal region of the embryo (“dorsal folds”, arrowheads in Fig. 1A). It is important to note that these defects in cell division and morphogenetic processes elsewhere in the embryo could also contribute to the reduced rate of germband extension observed in the *dlg1* RNAi embryos.

Interestingly, despite the strong disruption of junctional myosin upon deletion of Dlg1, the rate of AP junction shrinkage was largely normal in the *dlg1* RNAi embryos. Previous work has demonstrated that both junctional myosin and medioapical myosin contribute to AP junction shrinkage. Specifically, medioapical myosin is thought to drive junction shrinkage, whereas junctional myosin is thought to stabilize junctions after junction shrinkage (Rauzi et al., 2010). It is possible that the reduction in junctional myosin is largely compensated by the increase in medioapical myosin in the *dlg1* embryos. Alternatively, the residual junctional myosin in the *dlg1* embryos may be sufficient to stabilize the junction following medioapical myosin-mediated junction shrinkage. Another unexpected finding in our study was that loss of Dlg1 activity mainly affected the orientation, but not the average growth rate, of the newly formed DV junctions. Previous studies have shown that both local and global forces contribute to new junction extension (Collinet et al., 2015; Yu and Fernandez-Gonzalez, 2016). At the tissue level, invagination of the posterior midgut provides a pulling force that facilitates DV junction extension and helps orient newly formed junctions (Collinet et al., 2015). At the individual quartet level, manipulating apical actomyosin contractility locally can influence the rate and orientation of new interfaces as they form. In particular, ectopically increasing apical tension in the AP cell pair promotes DV junction extension, whereas ectopically increasing apical tension along the DV cell pair inhibits it (Yu and Fernandez-Gonzalez, 2016). In *dlg1* RNAi embryos, we observed ectopic apical constriction and elevated apical area fluctuation in the germband cells, suggesting that the apical tension is more variable over time and more heterogeneous across the tissue. Future studies analyzing the specific activities of medioapical myosin and apical tension in AP and DV cell pairs in the *dlg1* RNAi embryos may help to understand the specific defects we observed in T1 transitions.

Another question that remains to be addressed is the consequence of the ectopic accumulation of myosin along the lateral membrane in the *dlg1* RNAi embryos. A recent study shows that multicellular rosette formation, another mechanism that contributes to cell rearrangements during germband extension (Blankenship et al., 2006), occurs not only at the apical/subapical region of the epithelium, but also at the basolateral region (Sun et al., 2017). Interestingly, the formation of apical and basolateral rosettes is coordinated, yet involves distinct mechanisms (Sun et al., 2017). An interesting avenue for future research is to examine whether rosette formation is affected in the *dlg1* RNAi embryos and, in particular, whether ectopic lateral myosin accumulation impairs basolateral rosette formation.

The mechanism by which Dlg1 regulates myosin remains to be fully elucidated. Dlg1 may regulate myosin through direct or indirect manners. Moreover, regulation of myosin at the apical/subapical domain and at the lateral membrane may differ. *cyst* RNAi embryos exhibit myosin localization defects at the apical domain, but no observable myosin localization defects along the lateral membrane. In addition, knockdown of Cyst in the *dlg1* RNAi embryos inhibited ectopic accumulation of myosin along the lateral membrane, but not at the medioapical domain. These observations demonstrate that the myosin phenotypes at the apical/subapical domain and at the lateral domain can be decoupled. Our data suggest that Dlg1 regulate junctional myosin in a Cyst dependent manner. *cyst* deficient embryos show similar apical/junctional myosin phenotypes as *dlg1*, *scrib,* and *lgl* deficient embryos, including an increase in medioapical myosin and a decrease in junctional myosin (Silver et al., 2019 and this study). Furthermore, we show that Cyst-GFP is largely depleted from adherens junctions when Dlg1 is depleted. These observations suggest that Dlg1 promotes the activation of junctional myosin through the Cyst-Rho1-Rok pathway. The cause of ectopic medioapical myosin accumulation in *dlg1* RNAi embryos is unclear. Interestingly, loss of junctional myosin during germband extension has often been found associated with an increase in medioapical myosin, as demonstrated in the analyses of *dlg1*, *baz*, *crb*, and *cyst* mutant embryos (Silver et al., 2019; Simões et al., 2010; this study). Therefore, it is possible that these two myosin phenotypes are linked. Finally, our double RNAi experiment demonstrated that knockdown of Cyst in the *dlg1* RNAi background rescued the ectopic lateral myosin accumulation phenotype. These results suggest that ectopic accumulation of lateral myosin in *dlg1* RNAi embryos requires Cyst activity. The ectopic activation of lateral myosin could be a direct consequence of mis-localization of Cyst to the lateral membrane, where it activates myosin through Rho1 and Rok. Alternatively, Cyst could contribute to the lateral myosin phenotype indirectly by regulating other myosin regulators. Future experiments are required to further tease apart these possibilities.

Overall, the results presented in this work illustrate a novel molecular link between apical-basal polarity and cortical actomyosin contractility in epithelia. An important avenue of future research is to elucidate the molecular mechanism by which Dlg1 regulates the spatial organization of myosin and its upstream regulators. Through analysis of specific *dlg1* mutants, we observed that both the SH3 and GUK domains of Dlg1 are important for its function in regulating the spatial organization of myosin in the germband. Interestingly, previous studies have shown that there is an intramolecular association between the SH3 and GUK domains of Dlg1, and that this interaction is abolished by a mutation in the SH3 domain associated with the *dlg1*[m30] allele (McGee and Bredt, 1999). It is possible that this intramolecular interaction is required for the spatial regulation of myosin. Alternatively, the function of Dlg1 may be dependent on its ability to simultaneously bind multiple partners through its SH3 and GUK domains. Future experiments elucidating the molecular activity of Dlg1 involved in myosin regulation will further our understanding of how apical-basal polarity determinants control polarized subcellular organization and how such regulation influences the mechanics of tissue remodeling in epithelial morphogenesis.

## Materials and Methods

### Fly stocks and genetics

Sqh-GFP is described in (Royou et al., 2002). Sqh-mCherry is described in (Martin et al., 2009). UtrophinABD-Venus is described in (Figard and Sokac, 2011). The following RNAi lines were obtained from the Bloomington *Drosophila* Stock Center: *dlg1* TRiP (36771, 33620) (Fuentes and He, 2022), *lgl* TRiP (35773, 38989) (Fuentes and He, 2022), *scrib* TRiP (38199) (Fuentes and He, 2022), *baz* TRiP (39072) (Takeda et al., 2018), *cno* TRiP (33367) (Bonello et al., 2018), *cyst* TRiP (38292, 41579) (Garcia De Las Bayonas et al., 2019; Silver et al., 2019), *y* TRiP (64527). The TRiP lines were crossed to a maternal GAL4 driver line, Maternal-Tubulin-Gal4 67.15 (“Mat67; Mat15”) (Hunter and Wieschaus, 2000), for expression of shRNA during oogenesis (see below for details). AniRBD-GFP is described in (Munjal et al., 2015). Cyst-GFP is described in (Garcia De Las Bayonas et al., 2019). RhoGEF2-GFP (76260) (Nakamura et al., 2017) and Rok-GFP (52289) (Abreu-Blanco et al., 2014) lines were obtained from the Bloomington *Drosophila* Stock Center. The following *dlg1* temperature sensitive mutant lines were obtained from the Bloomington *Drosophila* Stock Center: *dlg1^2^*/FM7a (36278) and *dlg1^5^*/FM7a (36280). The following *dlg1* mutant lines were a gift from the Bilder lab: *dlg1^m52^*, *dlg1^m30^*, *dlg1^v59^*, *dlg1^IP20^*. An *ovo^D^* FRT19A (23880) line obtained from the Bloomington *Drosophila* Stock Center was used to generate *dlg1* germline clones with the FLP-dominant female sterile (DFS) system (see below for details).

To examine myosin localization defects in embryos with maternal knockdown of specific candidate genes via RNA interference (“RNAi embryos”), female flies from specific TRiP lines were crossed to Mat67 Sqh-GFP; Mat15 Sqh-GFP/TM3 males to generate Mat67 Sqh-GFP/TRiP; Mat15 Sqh-GFP/+ or Mat67 Sqh-GFP/+; Mat15 Sqh-GFP/TRiP flies. The embryos derived from these flies were used for analysis. Control embryos derived from Mat67 Sqh-GFP/+; Mat5 Sqh-GFP/+ flies, were F1s from the cross between *y w f* females and Mat67 Sqh-GFP; Mat15 Sqh-GFP/TM3 males.

*dlg1* germline clones were generated with the FLP-DFS system and *ovo^D^* FRT19A. *dlg1* FRT19A/FM7; Sqh-GFP/cyo females were crossed to *ovo^D^* FRT19A hsFLPase/FM7 males. *ovo^D^* is a dominant female sterile mutation. FRT-mediated recombination was induced with heat shock at 37°C. Females producing *dlg1*/*dlg1* germline clones were crossed to FM7/Y; Sqh-GFP/+ males to generate embryos used for analysis. Crosses with the temperature sensitive *dlg1* alleles were kept at room temperature (∼ 22°C).

Similar crosses were made to examine the localization of Rok-GFP, Cyst-GFP, AniRBD-GFP and RhoGEF2-GFP in RNAi embryos. The maternal GAL4 lines used for these studies were (1) Mat67 Sqh-mCherry/Cyo; Mat15 Rok-GFP, (2) Mat67 Sqh-mCherry/Cyo; Mat15 Cyst-GFP, (3) Mat67 Sqh-mCherry/Cyo; Mat15 AniRBD-GFP, and (4) Mat67 Sqh-mCherry/Cyo; Mat15 RhoGEF2-GFP.

### Live imaging and image processing

All live imaging was performed at room temperature. Embryos were dechorionated in ∼40% bleach (∼ 3% sodium hypochlorite), rinsed thoroughly with water, transferred on a 35 mm MatTek glass-bottom dish (MatTek Corporation), and covered with water.

In order to examine the rate of posterior midgut movement during germband extension, live imaging of control and *dlg1* mutant embryos expressing UtrophinABD-Venus was performed with an Olympus FVMPE-RS multiphoton microscope, a 25×/1.05 numerical aperture water-immersion objective, and a 920 nm pulsed laser. A 1× zoom was used. Single plane images (1024 pixels by 1024 pixels, or 509 µm by 509 µm) at the midsagittal plane of the embryo were acquired in 1-minute intervals beginning at the onset of VF formation. The lateral pixel size is 0.50 μm. Embryos were mounted with the lateral side facing the objective, such that both dorsal and ventral sides of the embryo were captured when imaged at the midsagittal plane. Representative images were aligned by time based on T_pmg_ (when the posterior midgut was 20 µm away from the posterior-most end of the embryo). The position of the posterior midgut was manually tracked over time. The velocity of posterior midgut movement was calculated between 60 μm and 140 μm from the posterior pole over.

To observe cell intercalation and cell division during germband extension, live imaging of control and *dlg1* RNAi embryos expressing UtrophinABD-Venus was performed with an Olympus FVMPE-RS multiphoton microscope, a 25×/1.05 numerical aperture water-immersion objective, and a 920 nm pulsed laser. A 3× zoom was used. A region of interest (ROI: 512 pixels by 256 pixels, 170 μm by 85 μm) that encompassed the lateral side of the embryo was imaged. Stacks of 36 images taken at 1-μm steps were acquired beginning at the onset of VF formation in 1-minute intervals with a Galvano scanner. Embryos were mounted with the ventrolateral side facing the objective to capture the ventrolateral ectoderm. Movies were aligned by time based on the onset of rapid VF invagination (approximately 10 minutes after the onset of apical constriction in the ventral mesoderm), which we defined as time zero in our analyses. In order to better quantify differences in tissue movement between the control and *dlg1* RNAi embryos, we manually tracked 7 cells in three control and three *dlg1* RNAi embryos over the course of an hour after time zero. These cells were located near the dorsal-most region of the ROI and were evenly spaced along the width (AP axis) of the ROI at time zero. The onset of cell division was determined as the first time point in which mitotic rounding could be observed within the ROI, which was located at the ventrolateral region of the embryo.

To examine the localization of myosin and myosin activators in the ventrolateral ectoderm of the control and *dlg1* RNAi embryos, deep tissue live imaging of ventrally oriented control and *dlg1* RNAi embryos was performed with an Olympus FVMPE-RS multiphoton microscope, a 25×/1.05 numerical aperture water-immersion objective, and a 920 nm pulsed laser. A 3× zoom was used. A region of interest (ROI: 512 pixels by 128 pixels, 170 μm by 42 μm) that encompassed the middle region along the AP axis was imaged. Stacks of 101 images taken at 1-μm steps were acquired beginning at the onset of VF formation in 2-minute intervals with a Galvano scanner. Cross section views encompassing the entire depth of the ventral side of the embryo were generated in ImageJ by re-slicing, followed by the maximum projection of 129 slices (∼ 43 μm thick stack along the AP axis). Flattened *en face* views of the embryo were generated by using a custom MATLAB script that fits the cross-section view of the embryo to a circle, and then flattens the *en face* views at a defined apical-basal depth from the surface. Embryos were aligned by time based on the onset of rapid VF invagination (approximately 10 minutes after the onset of apical constriction in the ventral mesoderm).

### T1 transition analysis

Ectodermal cells were segmented and manually corrected with Exploratory Development Geometry Explorer (EDGE), a MATLAB-based program for cell shape analysis (Gelbart et al., 2012). Three control embryos and four *dlg1* RNAi embryos were analyzed. For each embryo, a single focal plane near the apical surface of the ventrolateral ectoderm was selected for analysis. For different embryos, the size of the analyzed region ranged from 200 pixels by 100 pixels (67 μm by 33 μm) to 240 pixels by 120 pixels (80 μm by 40 μm). The duration of analyzed movies ranged from 30 to 36 minutes, starting from T ≈ 10 minutes. Since we focused on cell shape change and cell neighbor exchange in this analysis, we generated movies that were compensated for global tissue flow using a custom MATLAB script, before the cells were segmented and tracked with EDGE. We then manually identified individual quartets that would undergo T1 transitions from the segmented movies. 56 quartets from 3 control embryos and 63 quartets from 4 *dlg1* RNAi embryos were selected for analysis. The quartet was defined to be at the T1 configuration if the AP cell pair shared a common interface. This interface was called the AP junction. Conversely, the quartet was defined to be at the T3 configuration if the DV cell pair shared a common interface. This interface was called the DV junction. Finally, the quartet was defined to be at the T2 configuration if all the four cells shared a common vertex. The first T2 configuration was defined as the end of persistent AP junction shrinkage, whereas the last T2 configuration was defined as the beginning of persistent new DV junction extension. The duration of T2 was defined as the time span between the first and the last T2 configurations (Fig. 2E). The length and angle of the AP junction and DV junction for each quartet was measured and plotted over time (Fig. 2D). The angle of the junctions fell within 0° and 90°, with 0° being parallel with the AP axis and 90° being perpendicular to the AP axis (horizontal and vertical in the acquired images, respectively). To determine the average length and angle change during AP junction shrinkage, quartets were aligned based on the first T2 (Fig. 2F). To determine the average length and angle change during persistent new DV junction extension, quartets were aligned based on the last T2 (Fig. 2H). The rate of AP junction shrinkage was defined as the average of AP junction length reduction during an 8-minute interval immediately before the first T2 (Fig. 2G). The rate of DV junction extension was defined as the average DV junction length increase during an 8-minute interval immediately after the last T2 (Fig. 2I). To evaluate the impact of variation in the duration of T2 on new junction formation, average DV junction length over time was calculated after aligning quartets based on the first T2 (Fig. S3B, C).

To analyze the apical cell area change, the apical area of the four cells within each quartet was measured during a 20-minute interval, ranging from T = 11 – 30 minutes to T = 21 – 40 minutes for different quartets (Fig. 3B). These quartets were then aligned to the beginning of the 20-minute interval to determine the average quartet behavior (Fig. 3C-G). To demonstrate the variation in individual cell area within a quartet (Fig. 3C), cells within the same quartet were sorted based on their net area change within the 20-minute time interval and were placed into the following four categories: the cell with greatest negative area change, the cell with 2^nd^ greatest negative area change, the cell with 3^rd^ greatest negative area change, and the cell with least negative area change. Then, the area-over-time data for cells within each category were averaged between quartets from the same genotype. To further demonstrate cell area variation within a quartet, the standard deviation of apical area (Fig. 3E), or apical area change (Δarea, Fig. 3F), of four cells within the same quartet was measured. To evaluate cell area fluctuation over time (Fig. 3G), the area-over-time curve was first fit into a third-degree polynomial, which was used to de-trend the original curve. For each quarter, an area fluctuation index was calculated using the following formula:

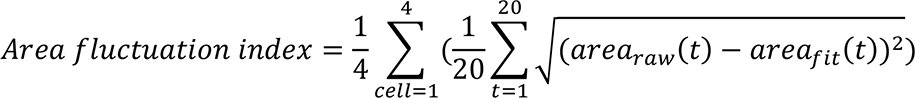

### Quantitative analysis of junctional and lateral GFP signals

To compare junctional enrichment of Sqh-GFP, Rok-GFP, Cyst-GFP, and AniRBD-GFP between different genotypes, we defined the “junctional signal index” based on the repeated “peak-valley-peak” pattern of intensity distribution across multiple AP cell boundaries observed in wildtype embryos (Fig. S4A, B). First, the spatial distribution of signal intensity along the AP axis for every DV position was extracted from the flattened *en face* view images, and the resulting signal was averaged every 8 pixels along the DV axis. Next, the discrete Fourier transform of the signal was calculated using the fft function in MATLAB with a sampling frequency of 18 pixels (6 μm -approximately one cell diameter), and a single-sided amplitude spectrum was created. We then defined the peak and background signal amplitude. The peak amplitude was defined as the maximal amplitude within the spectrum range of [0.70 - 1.27]. The background amplitude was defined as the average amplitude within the two flanking regions of the peak frequency ([0.42 - 0.56] and [1.41 - 1.55], respectively). Finally, the junctional signal index was calculated using the following formula:

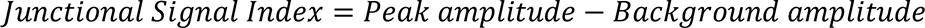

To compare the enrichment of Sqh-GFP, Rok-GFP, and Cyst-GFP at the lateral membranes between different genotypes, we defined the “lateral signal index” as described below (Fig. S4C). First, a region of interest (ROI) that covered the lateral region of the germband ectoderm at one side of the embryo was specified. Next, the mean pixel intensity within the ROI was determined from the maximum projection cross-section views at 0 minutes and 12 minutes. Finally, the lateral signal index was calculated using the following formula:

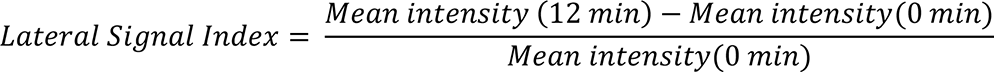

To compare the enrichment of AniRBD-GFP at the lateral membranes between control and *dlg1* RNAi embryos, we defined the lateral signal index using a different approach due to the lack of data at T = 0 min for most of the AniRBD-GFP movies acquired (Fig. S4C’). First, a region of interest (ROI) that covered the lateral region of the germband ectoderm at one side of the embryo and a background ROI (ROI_bg_) that covered the upper half of the ventral furrow was specified. Next, the mean pixel intensity within the ROI and ROI_bg_ was determined from the maximum projection cross-section views at 12 minutes. Finally, the lateral signal index was calculated using the following formula:

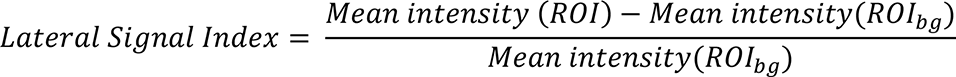

### Statistical Analysis

All quantification and data plotting were performed using MATLAB (MathWorks), R, or Microsoft Excel unless otherwise stated. Sample sizes for the presented data and methods for statistical comparisons can be found in figures or figure legends. Student t-test or Mann-Whitney U-test were used for all statistical analyses presented in this work. *p* values were calculated using the MATLAB ttest2 function and ranksum function, respectively.

## Author contributions

Conceptualization: B.H., M.A.F.; Methodology: B.H., M.A.F.; Software: B.H., M.A.F.; Validation: B.H., M.A.F.; Formal analysis: B.H., M.A.F., H.N.P; Investigation: M.A.F., H.N.P; Resources: B.H.; Data Curation: M.A.F.; Writing – original draft preparation: M.A.F.; Writing-review and editing: B.H., M.A.F., H.N.P; Visualization: B.H., M.A.F.; Supervision: B.H.; Project administration: B.H.; Funding acquisition: B.H.

## Acknowledgements

We thank A. Lavanway for critical help with imaging and research support in general. We thank D. Bilder and T. Lecuit for sharing reagents. We thank E. Griffin and J. Moseley for their valuable feedback on our manuscript. We thank M. Peifer for helpful discussion. We thank the B.H. lab members for constructive comments and discussion. We thank the Bloomington *Drosophila* Stock Center (NIH P40OD018537) for providing reagents used in this work. This research is supported by NIGMS ESI-MIRA R35GM128745 and American Cancer Society Research Grant #IRG-16-191-33 awarded to B.H, and the GAANN fellowship and the Ryan fellowship awarded to M.A.F.

The authors declare no further competing financial interests.

