## Supplementary Figures and Movie Legends for "Dlg1 regulates subcellular distribution of non-muscle myosin II during *Drosophila* germband extension"

#### Supplementary Materials

**Supplementary Figures 1 – 9**

**Supplementary Movie 1**

#### Supplementary Figures and Figure Legends

### Supplementary Figure 1

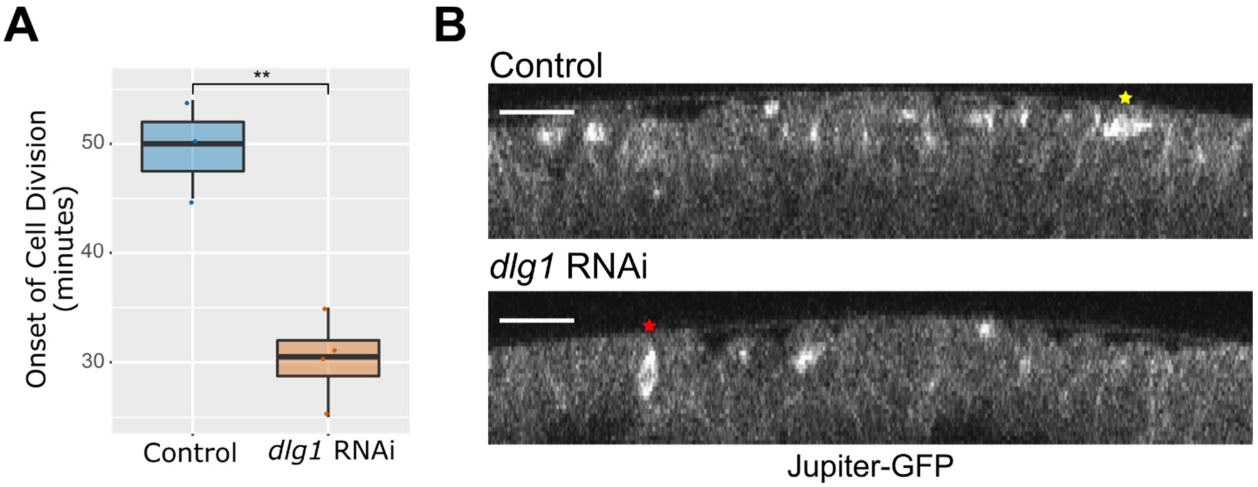


**Supplementary Figure 1. Mitosis defects in *dlg1* RNAi embryos**

**(A)** Quantification of cell division onset in the ventrolateral region of control (n = 3) and *dlg1* RNAi (n = 4) embryos. Unpaired, two-tailed student t-test was used for statistical analysis. **: $p \leq 0.01$. **(B)** *dlg1* RNAi embryos exhibit defects in the orientation of cell division. Movie stills showing the cross-section view of a representative control and *dlg1* RNAi embryo expressing the microtubule marker, Jupiter-GFP, during germband extension. In the control embryo, the yellow star indicates a cell undergoing proper cell division along the horizontal axis. In the *dlg1* RNAi embryo, the red star indicates a cell undergoing improper cell division along the vertical axis. Scale bars: 50 µm.

### Supplementary Figure 2

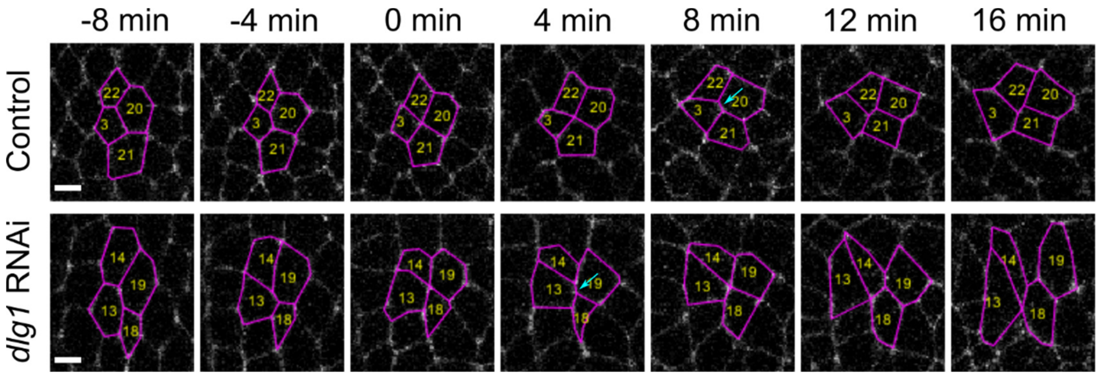


**Supplementary Figure 2. Prolonged T2 configurations can be observed in both control and *dlg1* RNAi embryos**

Examples of individual control and *dlg1* RNAi quartets undergoing a T1 transition followed by a prolonged T2 transition. Throughout the duration of the T2 transition several temporary T2-to-T1 reversals occur (arrows). Scale bars: 5 µm.

34 **Supplementary Figure 3**

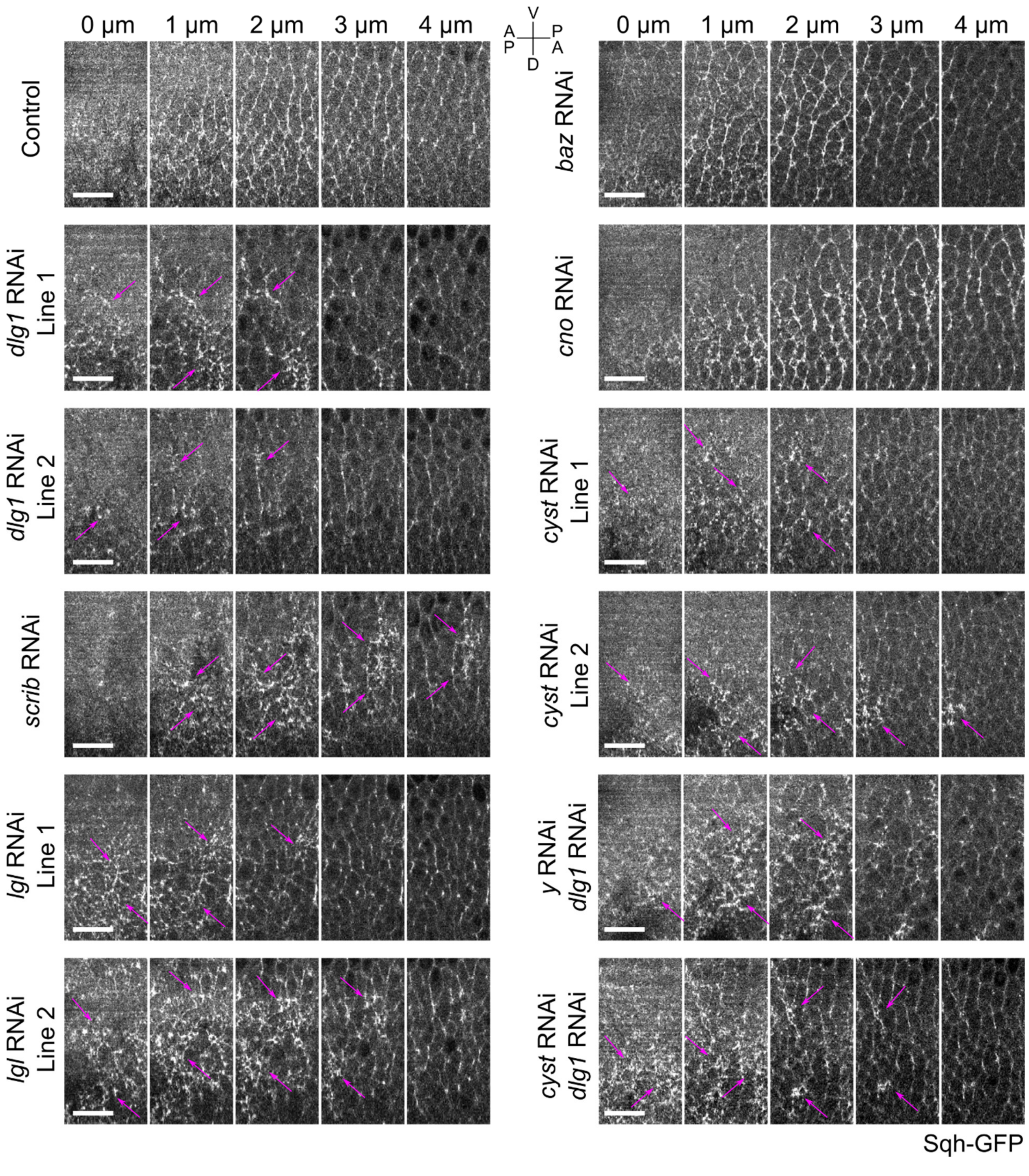


35

**Supplementary Figure 3. Apical myosin signal during early stages of germband extension in various RNAi backgrounds**

Montages showing the flattened *en face* views of a representative control embryo and representative embryos from various RNAi lines expressing the myosin marker, Sqh-GFP. 0 µm marks the apical surface. All images show embryos at T = 12 minutes after the onset of rapid VF invagination. Magenta arrows: ectopic medioapical myosin. Scale bars: 20 µm.

### Supplementary Figure 4

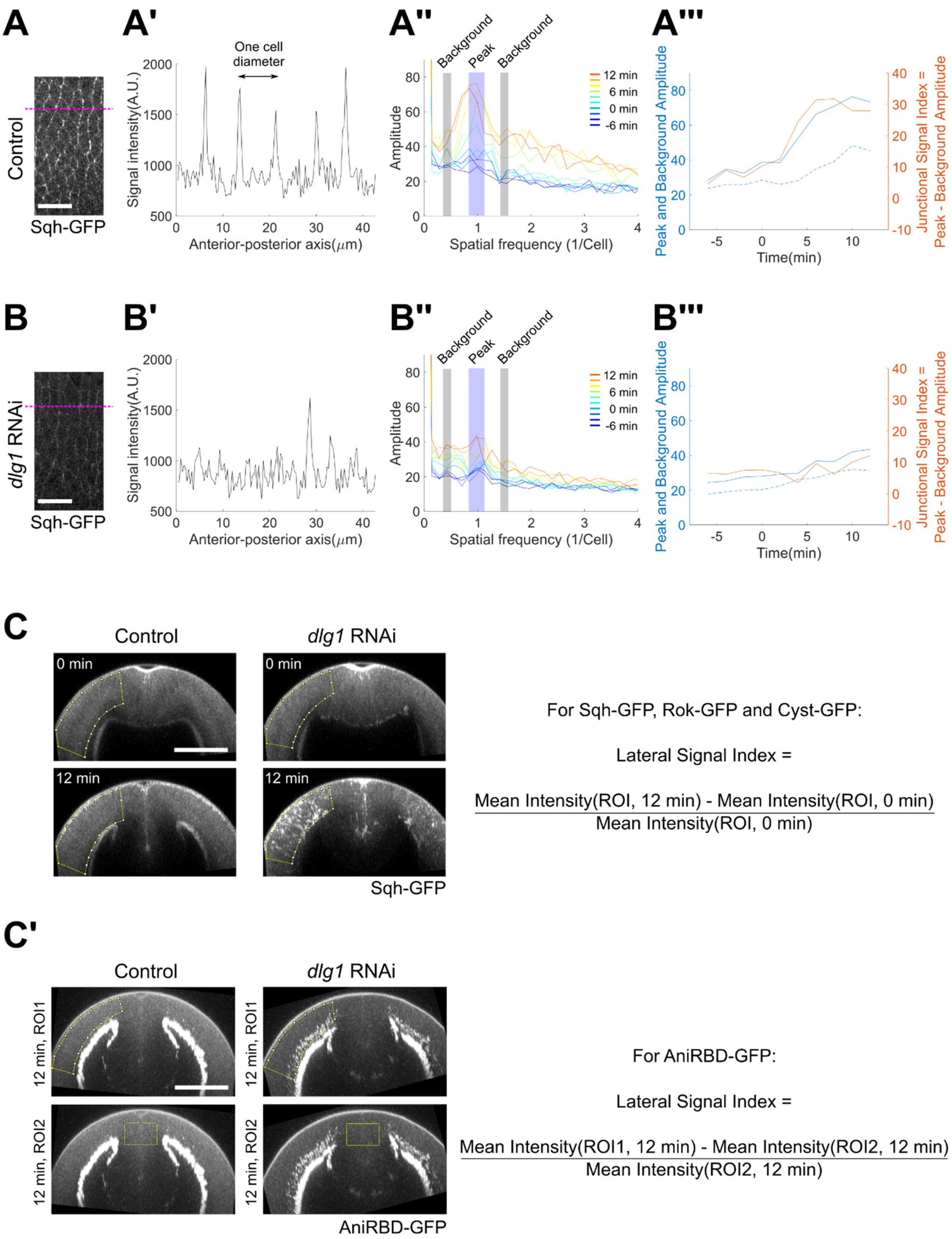

**Supplementary Figure 4. Analysis of GFP signal enrichment at the adherens junctions and the lateral membranes in the germband cells**

**(A, B)** Analysis of GFP signal enrichment at the adherens junctions. A representative control embryo (A) and a representative *dlg1* RNAi embryo (B) expressing Sqh-GFP are shown as examples. **(A’, B’)** Intensity distribution across the magenta lines in A and B. Peaks in A’ correspond to signal at the adherens junctions. **(A’’, B’’)** Fourier transform of the signals in A’ and B’. Results at different times after the onset of rapid VF invagination are plotted. In the control embryo, a prominent peak (magenta bar) is detected at the spatial frequency of one-cell diameter. This peak is much reduced in the *dlg1* RNAi embryo, corresponding to the reduced junctional signal in this background. **(A’’’, B’’’)** The peak amplitude, background amplitude, and junctional signal index over time. The background amplitude is defined as the average amplitude at the two background frequencies at two sides of the peak (gray bars in A’’ and B’’). The junctional signal index is defined as peak amplitude minus background amplitude. The same analysis is used to analyze Sqh-GFP, Rok-GFP, Cyst-GFP and AniRBD-GFP signals. **(C, C’)** Analysis of GFP signal enrichment at the lateral region of the germband cells. (C) Sqh-GFP, Rok-GFP and Cyst-GFP. (C’’) AniRBD-GFP. Note that a different formula is used for analyzing AniRBD-GFP due to the lack of data at T = 0 minute for most of the AniRBD-GFP movies acquired.

62 # Supplementary Figure 5

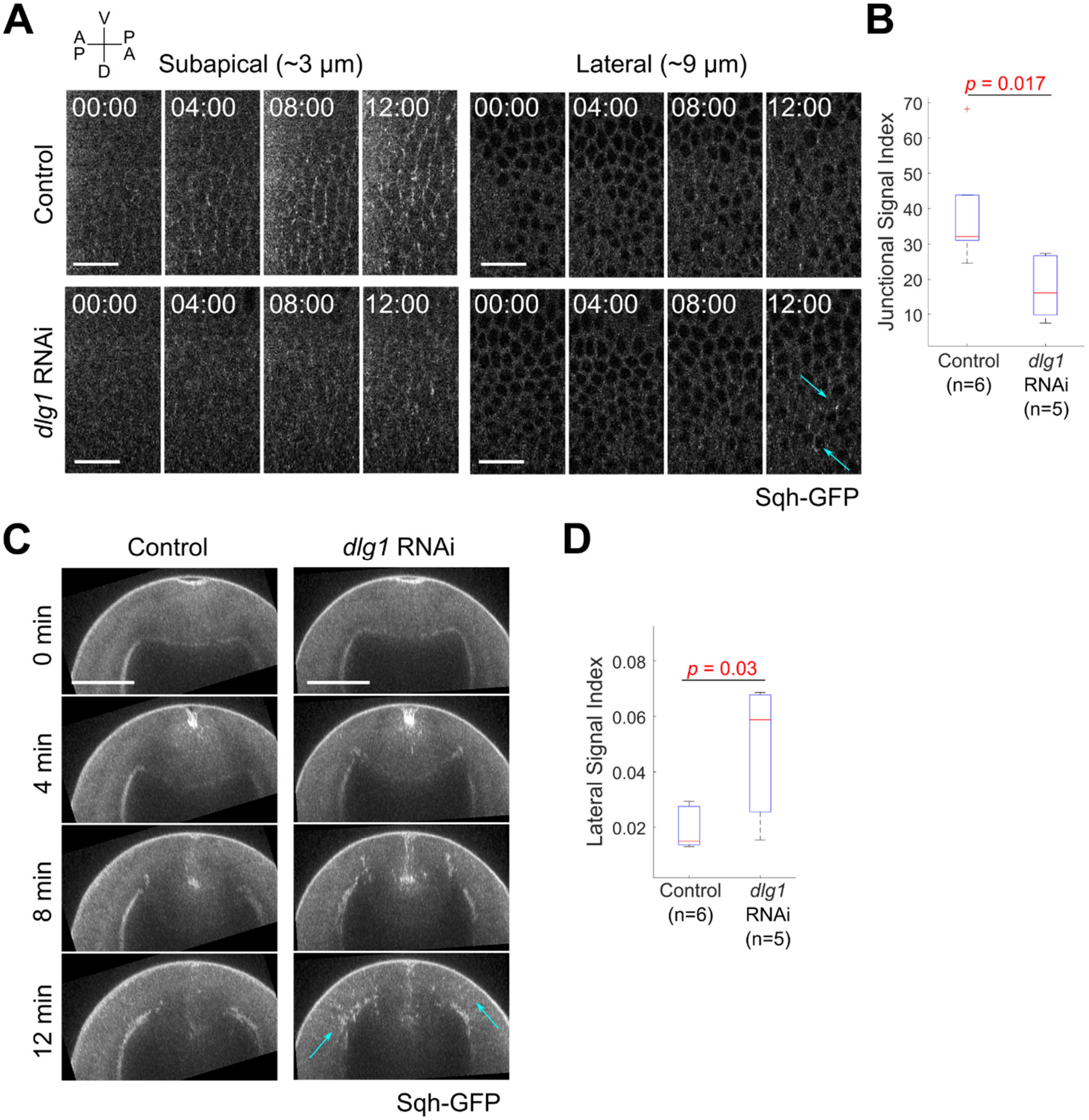


63

**Supplementary Figure 5. The *dlg1* RNAi myosin phenotypes visualized with one copy of Sqh-GFP**

**(A)** Movie stills showing the flattened *en face* view of a representative control and *dlg1* RNAi embryo expressing the myosin marker, Sqh-GFP, at depths of 3 µm (sub-apical region, left panel) and 9 µm (lateral region, right panel) from the surface of the embryo. T = 0 minutes indicates the onset of VF invagination. Cyan arrows: ectopic lateral myosin. Scale bars: 20 µm. **(B)** Junctional signal index of Sqh-GFP indicating the degree of myosin enrichment at the adherens junctions (Methods; Supplementary Figure 5). **(C)** Movie stills showing maximum projection cross-section views of the representative control and *dlg1* RNAi embryos shown in A. T = 0 minutes indicates the onset of VF invagination. Cyan arrows: ectopic lateral myosin. Scale bars: 50 µm. **(D)** Lateral signal index of Sqh-GFP indicating the degree of myosin enrichment at the lateral region of the germband epithelium (Methods; Supplementary Figure 5). Two-sided Mann-Whitney U-test was used for statistical comparison.

### Supplementary Figure 6

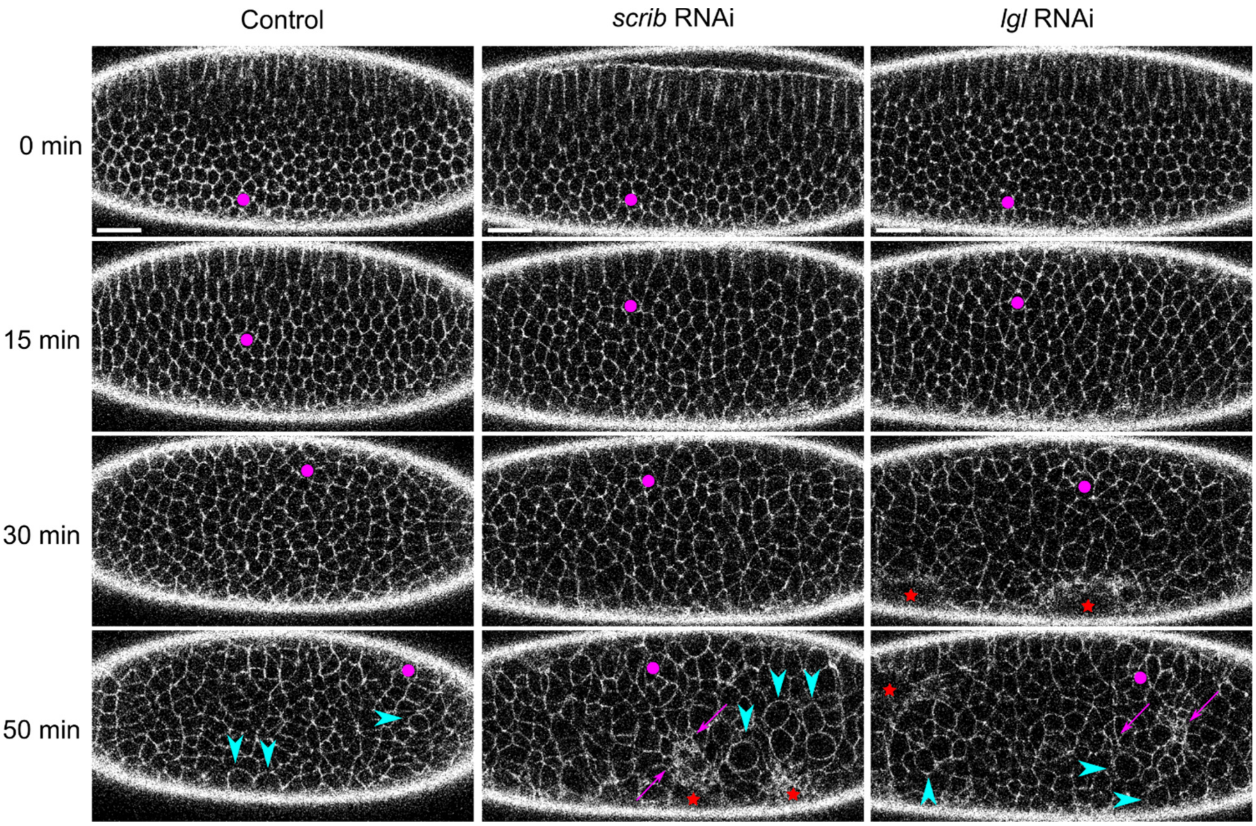

**Supplementary Figure 6. *scrib* and *lgl* RNAi embryos exhibit ectopic apical constriction during germband extension**

Movie stills showing the sub-apical plane of a ventrolaterally oriented representative control, *scrib* RNAi, and *lgl* RNAi embryo expressing UtrophinABD-Venus during germband extension. The ventral side is facing up. T = 0 minutes corresponds to the onset of rapid VF invagination. Magenta dots indicate individual cells that have been tracked over time. Both *scrib* and *lgl* RNAi embryos exhibit ectopic apical constriction (magenta arrows) like the *dlg1* RNAi embryos. Red stars indicate the formation of apical indentations associated with ectopic apical constriction in the RNAi embryos. Cyan arrowheads indicate cells that round up before cell division. Scale bars: 20 μm.

91 **Supplementary Figure 7**

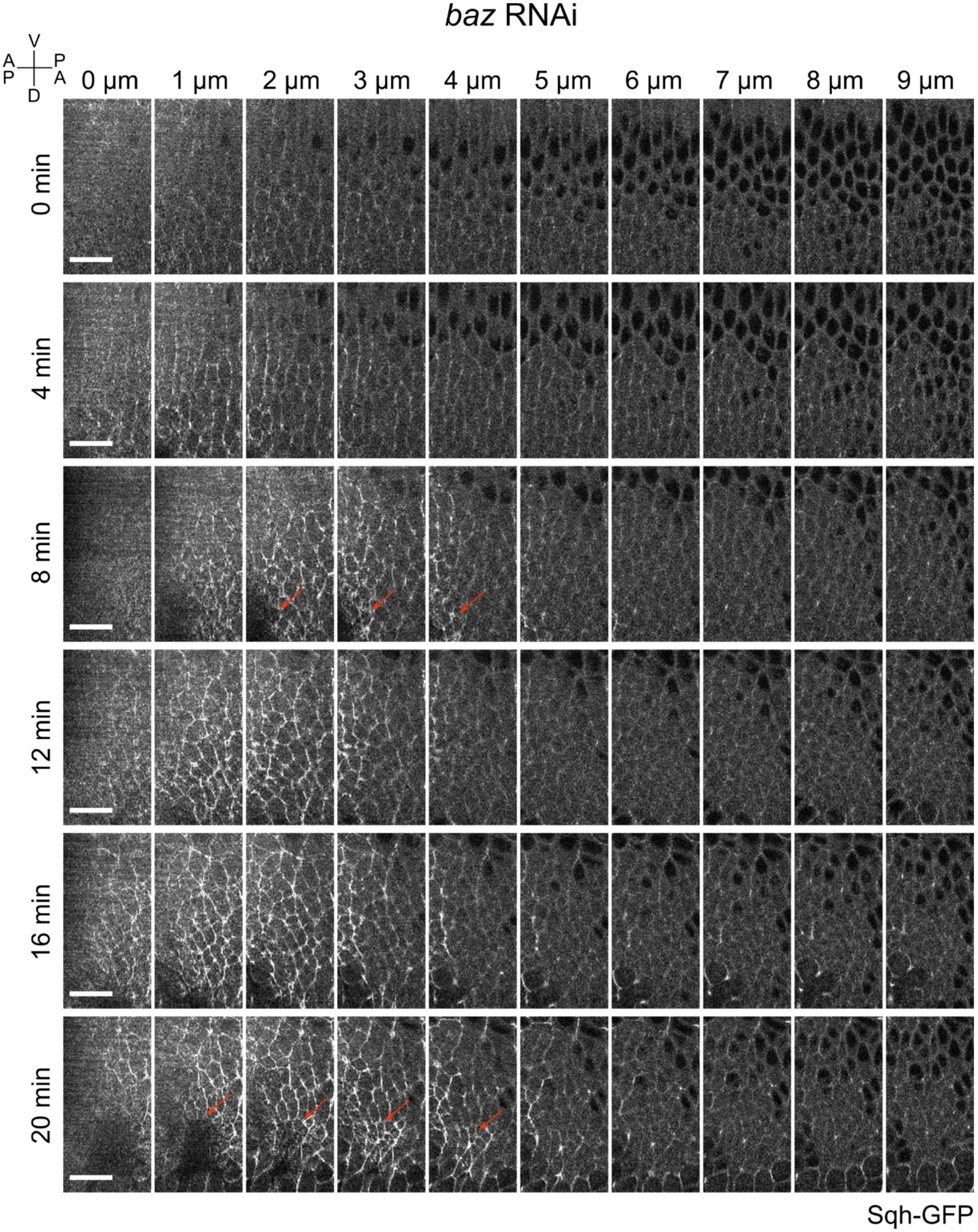


92

**Supplementary Figure 7. Ectopic fold/pit formation in *baz* RNAi embryos**

Montages showing the flattened *en face* views of a representative *baz* RNAi embryo expressing the myosin marker, Sqh-GFP. 0 μm marks the apical surface. Orange arrows: ectopic folds/pits. Note that only a moderate increase of medioapical myosin was observed in the regions of folds/pits. Scale bars: 20 μm.

98 **Supplementary Figure 8**

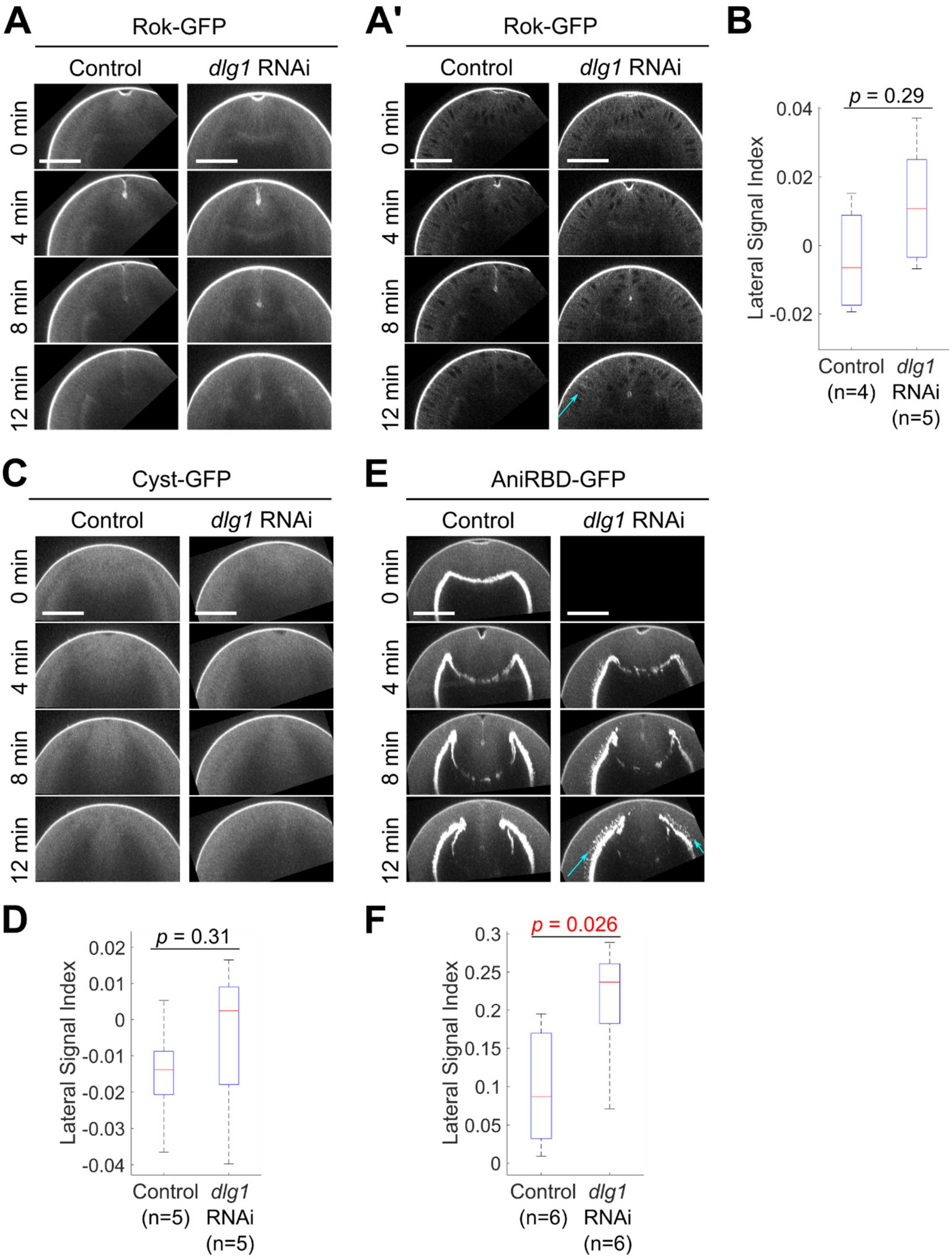


99

**Supplementary Figure 8. Cross-section views of representative control and *dlg1* RNAi embryos expressing Rok-GFP, Cyst-GFP, and AniRBD-GFP at early stages of germband extension**

**(A, C, E)** Movie stills showing maximum projection cross-section views of representative control and *dlg1* RNAi embryos expressing Rok-GFP (A), Cyst-GFP (C), and AniRBD-GFP (E). (A') shows projection of Rok-GFP over a single cell diameter where ectopic accumulation of Rok-GFP at the lateral membrane is occasionally observed (cyan arrow). T = 0 minutes indicates the onset of rapid VF invagination. The black images in (E) indicate data not acquired. Scale bars: 50 μm. **(B, D, F)** Lateral signal index of Rok-GFP (B), Cyst-GFP (D) and AniRBD-GFP (F) indicating the degree of signal enrichment at the lateral region of the germband epithelium (Methods; Supplementary Figure 5). Two-sided Mann-Whitney U-test was used for statistical comparison.

### Supplementary Figure 9

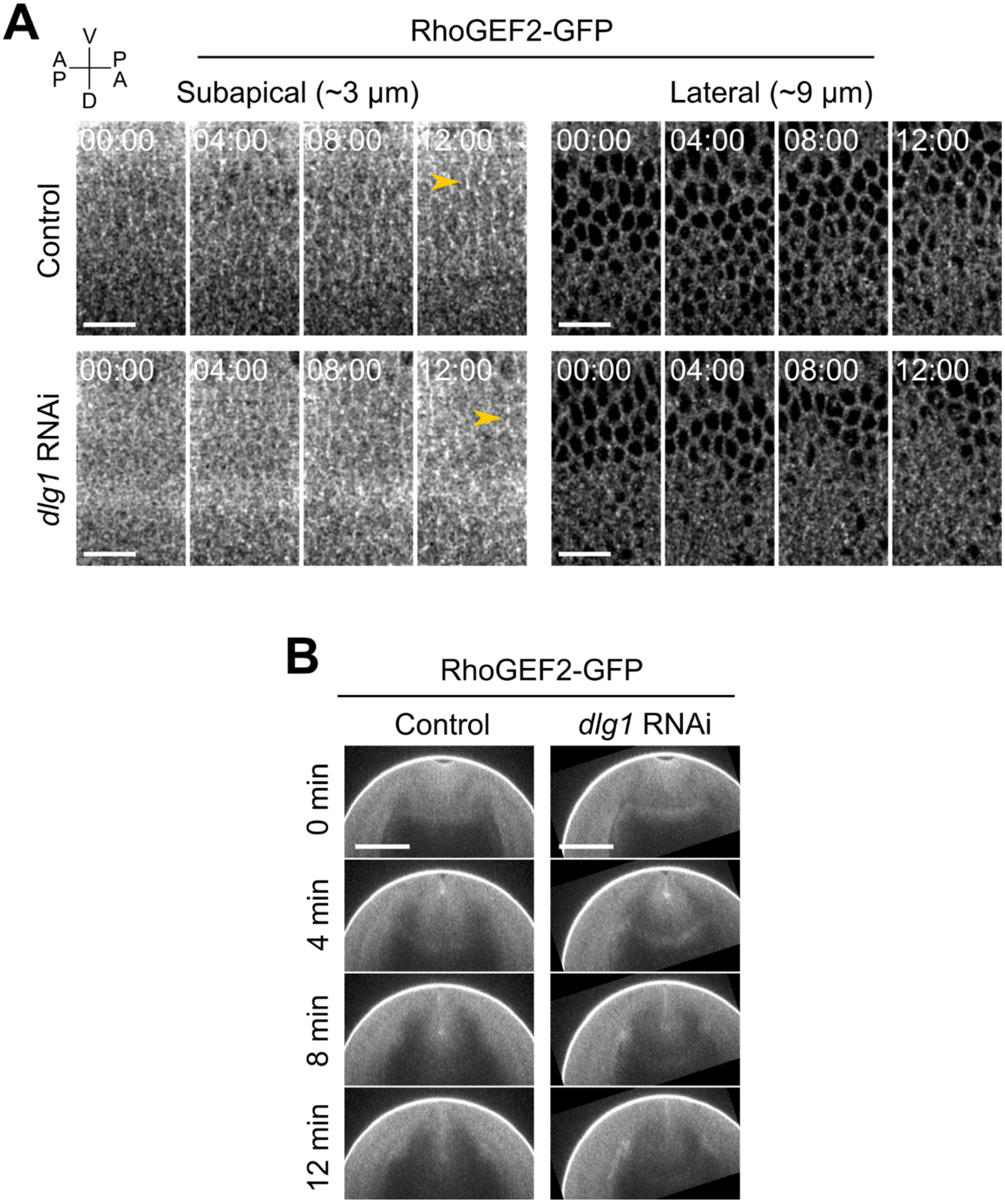

**Supplementary Figure 9. Knockdown of Dlg1 does not substantially impact the localization of RhoGEF2 at early stages of germband extension**

**(A)** Movie stills showing the flattened *en face* view of a representative control and *dlg1* RNAi embryo expressing RhoGEF2-GFP at depths of 3 µm (sub-apical region, left panel) and 9 µm (lateral region, right panel) from the surface of the embryo. A gaussian blur filter was applied to images shown in A. T = 0 minutes indicates the onset of VF invagination. Orange arrowheads: AP junctions. Scale bars: 20 µm. **(B)** Movie stills showing maximum projection cross-section views of the representative control and *dlg1* RNAi embryos shown in A. T = 0 minutes indicates the onset of rapid VF invagination. Scale bars: 50 µm.

#### Supplementary Movie

**Movie 1: Early stage of germband extension in control and *dlg1* RNAi embryos.**

Composite movie showing the subapical plane of a ventro-laterally oriented representative control and *dlg1* RNAi embryo expressing UtrophinABD-Venus during germband extension. The ventral side is facing up. T = 0 minutes corresponds to the onset of rapid VF invagination. Colored dots indicate the individual cells that have been tracked over time.
